# The phage Φ13-encoded transcriptional regulator Ltr controls phage assembly in Staphylococcus aureus

**DOI:** 10.1101/2025.11.28.691083

**Authors:** Ronja Dobritz, Marcel Bäcker, Carina Rohmer, Natalya Korn, Vittoria Bisanzio, Christiane Wolz

## Abstract

Temperate phages play a central role in evolution and pathogenicity of *Staphylococcus aureus*. Sa3int phages, in particular, contribute highly human-specific virulence factors that promote immune evasion and survival within the host. The reversible excision of these phages which occurs without phage production and bacterial lysis allows the simultaneous expression of phage virulence genes and the *hlb* gene where they usually integrate. However, the regulatory mechanisms controlling phage assembly and the cross-talk with host factors remain poorly understood. In this study, we analyzed the regulatory mechanism controlling late gene transcription in Sa3int phage Φ13. We identified a functional promoter, P_23,_ located upstream of the late phage genes that control DNA processing and packaging, capsid assembly, bacterial lysis and immune evasion. *SAOUHSC_02200*, the gene located upstream of P_23_, encodes for a late transcriptional regulator (Ltr). Mutating the P_23_ TATA-box or the *ltr* gene abolished P_23_ activity and formation of mature intact phage particles, thus confirming the role of Ltr in regulating P_23_ activity. Four direct repeats upstream of the P_23_ transcriptional start site were identified as potential Ltr binding sites. RT-qPCR analysis confirmed that Ltr-dependent P_23_ activation is essential for expression of late genes and the subsequent propagation of Φ13. Furthermore, comparative analysis of P_23_ activity and *ltr* expression in different host strain backgrounds revealed strain-specific differences that appear to depend on the alternative sigma factor SigB and its downstream effector SpoVG. These findings establish Ltr as the major regulator of late gene expression in Φ13 and reveal bacterial host factors that control successful phage assembly, and bacterial lysis.

**Importance:** The dynamic integration and excision of highly prevalent Sa3int phages in *Staphylococcus aureus* is considered a regulatory switch that enables bacterial adaptation to specific niches. These phages carry several human-specific virulence genes and integrate into the *hlb* virulence gene. It was assumed that they undergo a mechanism termed ’active lysogeny’, which allows the phages to be excised reversibly without phage production or bacterial lysis. Here, we have identified a new phage-encoded ’late transcriptional regulator’ (Ltr) that controls the expression of all late phage genes and thus phage assembly and lysis. We found that Ltr activity is regulated by the alternative sigma factor B and its downstream effector SpoVG. Restriction of SpoVG, and consequently phage assembly, likely contributes to the maintenance of Sa3int phages, even under phage-inducing conditions. This may be relevant in certain infectious conditions where both the phage-encoded virulence genes and the gene that is usually interrupted by the phage are required for infectivity.

## Introduction

*Staphylococcus aureus* is a major opportunistic pathogen that asymptomatically colonizes the nose of up to 30% of the human population and can cause severe infectious diseases. The virulence of *S. aureus* can be influenced by mobile genetic elements, such as pathogenicity islands, plasmids and bacteriophages that encode accessory virulence factors (1).

All known temperate staphylococcal phages belong to the family of *Siphoviridae*, which are characterized by an icosahedral head, a long, non-contractile tail and dsDNA. The phage genomes are organized into functional modules for lysogeny, replication, DNA packaging, structural genes encoding for head and tail proteins and lysis (2–6). *S. aureus* can carry several prophages at the same time, some of those encode for various staphylococcal virulence factors (6, 7). Phages infecting *S. aureus* have been classified into seven major groups based on the encoded *integrase* gene (Sa-*int* type). The *int* type dictates the *attB* site for integration of the prophage into the bacterial genome (3).

With up to 96% abundance, Sa3int phages are the most prevalent temperate phages in human nasal isolates of *S. aureus* (3, 8). Phages of this group integrate into the *hlb* locus, disrupting the *hlb* gene encoding a sphingomyelinase (β-hemolysin) (3, 9). On the other hand, Sa3int phages carry a set of genes for human-specific virulence factors, including a staphylokinase, as well as chemotaxis- and complement inhibitors, on the so called Immune Evasion Cluster (10, 11).

Sa3int phages are repeatedly lost when *S. aureus* is transferred from humans to animals, confirming the importance of these phages for the adaptation of *S. aureus* to the human host. Interestingly, when animal-adapted strains were transmitted back to humans, they often re-acquire Sa3int phages (11, 12). It was also found that Sa3int phages are more prevalent in strains that colonize the human nose than in infectious strains (13).

Depending on the host environment, both the insertion and excision of prophages may confer a fitness advantage to the host bacterium. Recently, it was demonstrated that the integration of Sa3int prophages significantly reduced the uptake into human macrophages, thereby mediating escape from human innate immune cells (14). On the other hand, the excision of Sa3int phage Φ13 is a critical step in the pathogenesis of *S. aureus* during murine infections (15) and spontaneous Hlb-positive variants are selected during murine skin colonization (16) or infective endocarditis in rabbits (17). Sa3int phages may also undergo pseudo-lysogeny/active lysogeny a process during which a phage is temporally excised from the chromosome without forming intact phage particles (18–21). This is consistent with the observation that infection of *S. aureus* with Sa3int phages usually does not result in the lysis of the bacterial population (22). Through this process, bacteria can simultaneously activate phage virulence genes as well as the gene that is typically inactivated by phage integration (23).

Molecular cross-talk between Sa3int phages and their *S. aureus* host and regulatory processes controlling active lysogeny remain largely unknown. Previous studies revealed that prophage excision, replication and transfer is highly strain dependent (24, 25) Comparative RNA-Seq analysis of high (MW2c) and low transfer (8325–4) strains revealed significant differences in the expression of structural module genes (24). Thus, so far unidentified bacterial factors are crucial determinants for the phage life cycle.

Here, we investigated the transcriptional regulation of Sa3int phages to uncover the mechanisms that control phage assembly. We identified an essential promoter region within the phage genome that regulates late phage genes and discovered strain-specific differences in its activity. Moreover, we identified SAOUHSC_02200 as an activating regulator, with similarity to Family IV of the previously described late transcriptional regulators (Ltrs). Strain-specific differences in P_23_ activity implicated that the alternative sigma factor B (SigB) and its downstream effector SpoVG play a role in the regulation of P_23_ dependent phage gene expression and consequently phage assembly.

## Results

### The predicted promoter region P_23_ in the Φ13 genome is functional and exhibits strain-specific activity

In previous work, we identified 57 transcriptional start sites (TSS) across the Φ13 genome analyzed in low and high phage transfer strains (24). TSS23 became of interest, as it is located upstream of the structural module and precedes genes (*SAOUHSC_02198* to *SAOUHSC_02177*) that were significantly upregulated in the high-transfer strain MW2c. TSS23 is located in an intergenic region in between two genes of unknown function (upstream: *SAOUHSC_02200*; downstream: *SAOUHSC_02199*). Late phage genes including DNA-packaging genes (*endonuclease*, small (*terS*) and large terminase (*terL*)) and structural genes (*mcp*) are located downstream of TSS23. Sequence analysis of the region upstream of TSS23 revealed a putative TATA-box, suggesting the presence of a functional promoter. To analyze the activity and regulation of this putative promoter region (designated P_23_), we constructed reporter plasmids, covering 200 bp upstream and 50 bp downstream of the predicted TSS23 (Figure 1A). The P_23_ promoter region was fused to *yfp*, allowing fluorescence measurements to assess P_23_ activity.

**Figure 1:**
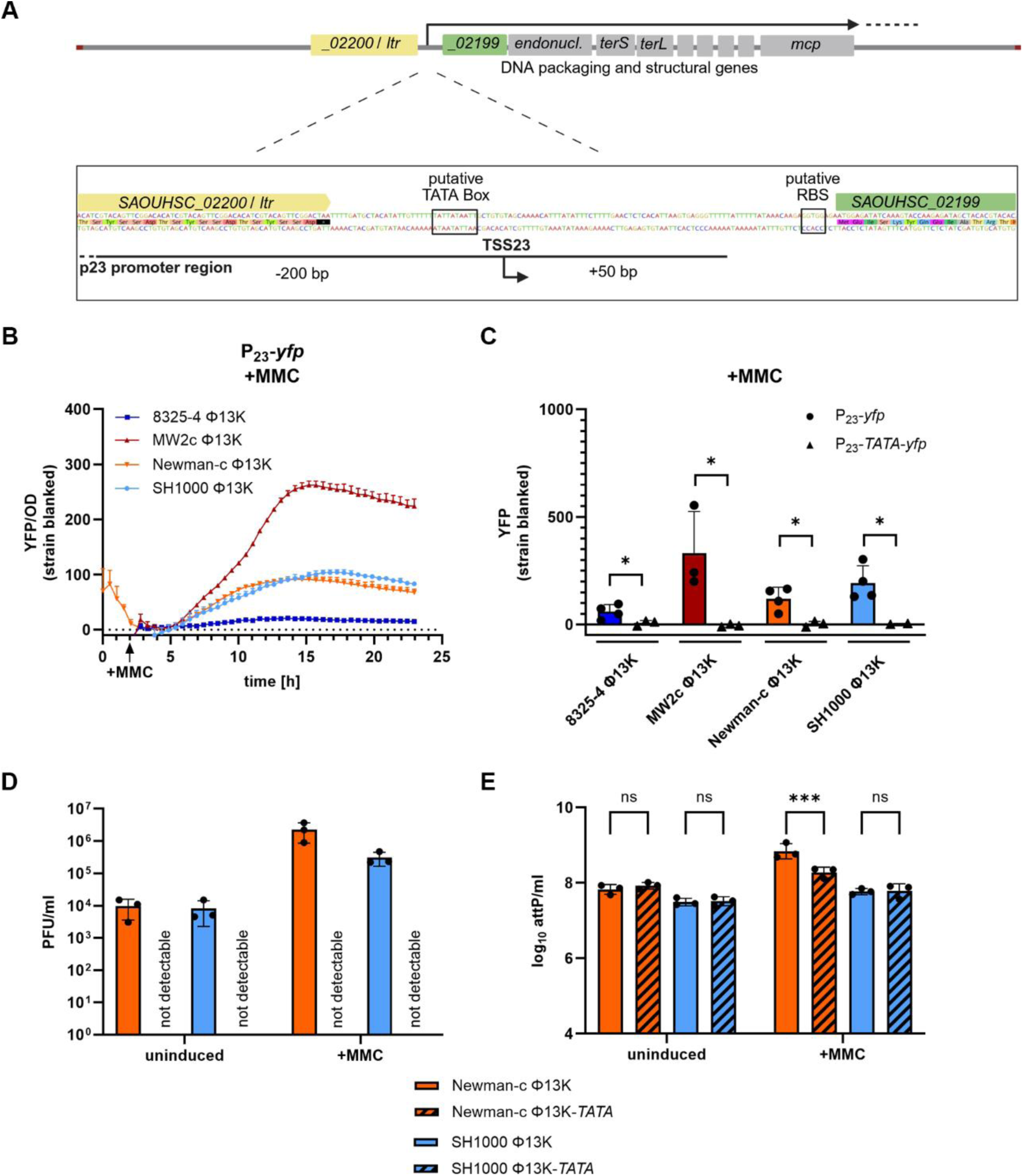
P_23_ is a functional and essential promoter of Φ13 with strain-specific activity. (A) Schematic depiction of P_23_ promoter region and transcriptional start site 23 (TSS23). (B) P_23_ promoter activity measurement over time after mitomycin C (MMC) addition. Single-lysogenic strains were grown to exponential growth phase and induced with a subinhibitory concentration of mitomycin C (300 ng ml^-1^). Optical density and fluorescence were measured for further 24 hours. Arbitrary units of fluorescence (YFP) are shown normalized to OD_600_ after substracting strain-specific background fluorescence. (C) Single time point measurement of P_23_ promoter activity without (P_23_-*yfp)* and with mutant TATA-Box (P_23_-*TATA*-*yfp*). Single-lysogenic strains were grown to exponential phase, induced with a subinhibitory concentration of MMC, and grown for additional 4 hours. Bacteria were harvested to OD_600_ of 2 and resuspended in PBS. Fluorescence was measured. Arbitrary units of fluorescence (YFP) are shown; strain-specific background fluorescence was subtracted. Data shown are mean ± SD (n ≥ 3). Statistical significance was determined by unpaired t-tests within strains. (D,E) Phage replication in strains carrying Φ13K (wild type) and Φ13K-*TATA* (TATA-box muation) under uninduced and induced (+MMC) conditions. Single-lysogenic Φ13K strains were induced with subinhibitory MMC in exponential phase and incubated for 60 min. Phage numbers were determined by plaque assay (D) and phage genome numbers by qPCR on the attachment site (*attP*) (E). Data shown are mean ± SD (n = 3). Statistical significance was determined by 2way ANOVA tests on log_10_ transformed data (***p-value < 0.001, *p-value < 0.05, ns > 0.05).

Promoter activity was measured for 24 h in four different Φ13K single-lysogenic *S. aureus* strains. Under uninduced conditions, P_23_ activity remained undetectable, indicating the promoter remains inactive in the lysogenic state (Supplemental Figure S1). Upon addition of subinhibitory concentrations of mitomycin C (MMC) to induce the bacterial SOS response and phage excision, strain-specific differences in P_23_ activity were obvious. The highest P_23_ activity was found in high-transfer strain MW2c Φ13K, while intermediate fluorescent signals were detected in Newman-c Φ13K and SH1000 Φ13K, and the lowest activity in 8325-4 Φ13K (Figure 1B). To assess strain-specific differences independently of growth effects, we measured P_23_ activity at single time-points with adjusted optical densities. Measurement at 4 hours after induction with MMC revealed the same strain-specific differences in P_23_ activity (Figure 1C).

To confirm promoter functionality and characteristics of P_23_, we replaced the putative TATA-box (position -15 to -5 relative to TSS23) with Cs and Gs (Figure 1A). The substitutions abolished P_23_ promoter activity, confirming that the region contains a functional TATA-box at the proposed position, which is required for promoter activity (Figure 1C, Supplemental Figure S1).

To investigate the role of P_23_ in phage replication and production, we mutated the TATA-Box in the P_23_ promoter region of the Φ13 genome. This resulted in a loss of phage particle production under uninduced and MMC induced conditions as determined by plaque assay (Figure 1D).

However, phage genome replication, quantified by qPCR still occurred in the TATA-Box mutant phage. Since no infectious phage particles were detectable, we assume that phage genomes are likely released into the supernatant through phage-independent mechanisms of bacterial cell lysis. Upon addition of MMC, no (SH1000) or only a slight increase (Newman-c, factor 2) in phage genome copy numbers was observed for the Φ13K-*TATA* phage mutant (Figure 1E). These results indicate that the phage loses its ability to produce intact mature phage particles through P_23_ disruption. However, phage genome replication was largely independent of P_23_ activity.

These findings confirm that the P_23_ promoter region is functional and essential for Φ13 propagation. Our data also indicates strain-specific differences in P_23_ activity potentially caused by host-dependent regulatory factors.

### *sigB* and *spoVG* dependent P_23_ regulation

*S. aureus* SH1000, which exhibited intermediate P_23_ activity, and *S. aureus* 8325-4, which showed the lowest activity, differ only by a defective *rsbU* gene in 8325-4. Since RsbU is responsible for the activation of SigB, 8325-4 is considered SigB-deficient. This was confirmed by analyzing the expression of *asp*, a prototypic SigB target gene (Figure 2A). This observation led us to investigate the involvement of SigB in P_23_ regulation.

**Figure 2:**
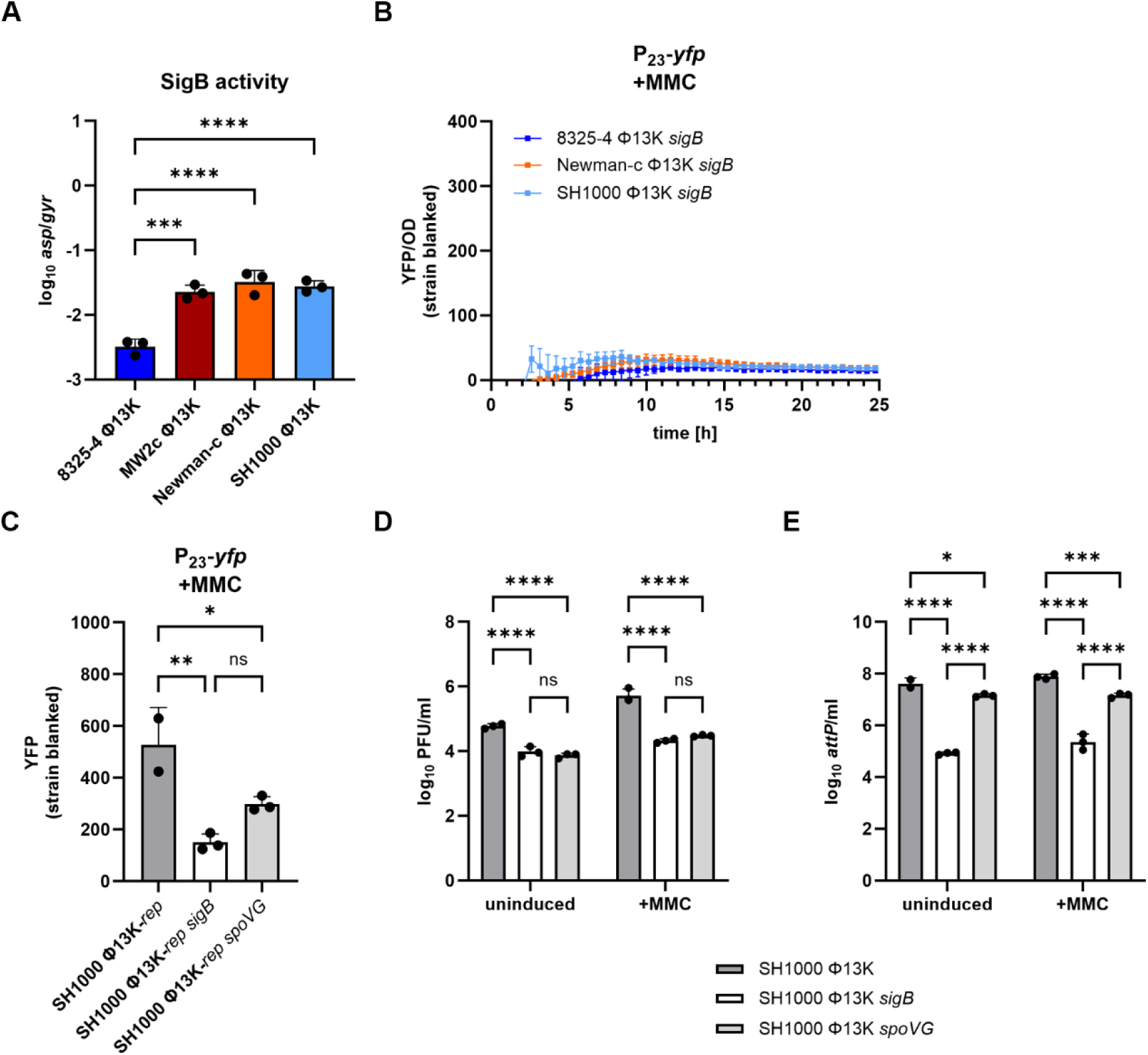
SigB and SpoVG impact P_23_ promoter activity and phage production. (A) Strain-specific SigB activity under induced conditions. Single-lysogenic strains were grown to exponential phase, induced with a subinhibitory concentration of mitomycin C (MMC) (300 ng ml^-1^), and grown for further 4 hours. Bacteria were harvested for RNA isolation and RT-qPCR was performed. The expression of *asp* was normalized to *gyrB* expression. Data shown are mean ± SD (n = 3). Statistical significance was determined by ordinary one-way ANOVA test on log_10_ transformed data. (B) P_23_ promoter activity in a *sigB*-deficient background over time after MMC addition. Single-lysogenic strains were grown to exponential phase and induced with a subinhibitory concentration of MMC. Optical density and fluorescence were measured for further 24 h. Arbitrary units of fluorescence (YFP) are shown normalized to OD_600_; strain-specific background fluorescence was subtracted. (C) Single time point measurement of P_23_ promoter activity in SH1000 (wild type), SH1000 *sigB* (*sigB*::*tet* deletion), and SH1000 *spoVG* (*spoVG*::*erm* deletion). Single-lysogenic strains were grown to exponential phase, induced with a subinhibitory concentration of MMC and grown for further 4 hours. Bacteria were harvested to OD_600_ of 2 and resuspended in PBS. Fluorescence was measured. Arbitrary units of fluorescence (YFP) are shown; strain-specific background fluorescence was subtracted. Data shown are mean ± SD. Statistical significance was determined by ordinary one-way ANOVA. (D,E) Phage replication in single-lysogenic SH1000 (wild type), SH1000 *sigB* (*sigB*::*tet* deletion) and SH1000 *spoVG* (*spoVG*::*erm* deletion) under uninduced and induced (+MMC) conditions. Single-lysogenic Φ13K strains were induced with subinhibitory MMC in exponential growth phase and incubated for 60 min. Phage numbers were determined by plaque assay (D) and phage genome numbers by qPCR on the attachment site (*attP*) (E). Data shown are mean ± SD (n = 3). Statistical significance was determined by 2way ANOVA tests on log_10_ transformed data. (****p-value < 0.0001, ***p-value < 0.001, **p-value < 0.01, *p-value < 0.05).

Deletion of the whole *sigB* operon (Δ*mazEFrsbUVWsigB*) in single-lysogenic strains resulted in a decrease of P_23_ activity to levels comparable to those of the natural SigB-deficient strain 8325-4 (Figure 2B). P_23_ activity after 4 hours of induction with MMC was fourfold lower in the *sigB* deleted SH1000 single-lysogen than in the wild type. Since P_23_ promoter region does not contain a SigB consensus motif, the SigB effect is likely mediated by one of its downstream effectors. Expression of the putative regulator *spoVG* is directly regulated through SigB and might serve as SigB effector. lndeed, mutation of *spoVG* resulted in a similar decrease in P_23_ promoter activity to that observed for the *sigB* mutation (Figure 2C). Deletion of either *sigB* or *spoVG* also resulted in significantly reduced phage particle numbers as quantified by plaque assay (Figure 2D). However, the number of phage genome copies decreased more profoundly in the *sigB* mutant than in the *spoVG* mutant indicating that SigB exerts some additional SpoVG-independent effects on phage replication (Figure 2E).

Overall, these results demonstrate that SigB and its downstream effector SpoVG directly or indirectly modulate P_23_ activity and subsequent phage propagation.

### A phage factor is required for P_23_ activity

We further analyzed whether phage factors are involved in P_23_ activity. Therefore, using the reporter plasmid with the wild type promoter region, we analyzed P_23_ activity in bacterial strains carrying various prophage mutants, as well as in a prophage-free background. Promoter activity was significantly increased in single-lysogens carrying either the P_23_-disrupted prophage mutant (Φ13K-*TATA*) or the replication-deficient prophage (Φ13K-*rep*) (Figure 3A). We speculate that the enhanced promoter activity may be the result of increased binding of a regulatory factor to the reporter plasmid in the absence of a functional target site in the prophage genome (for Φ13K-TATA) or due to a reduced number of genomic promoter copies in Φ13K-rep.

**Figure 3:**
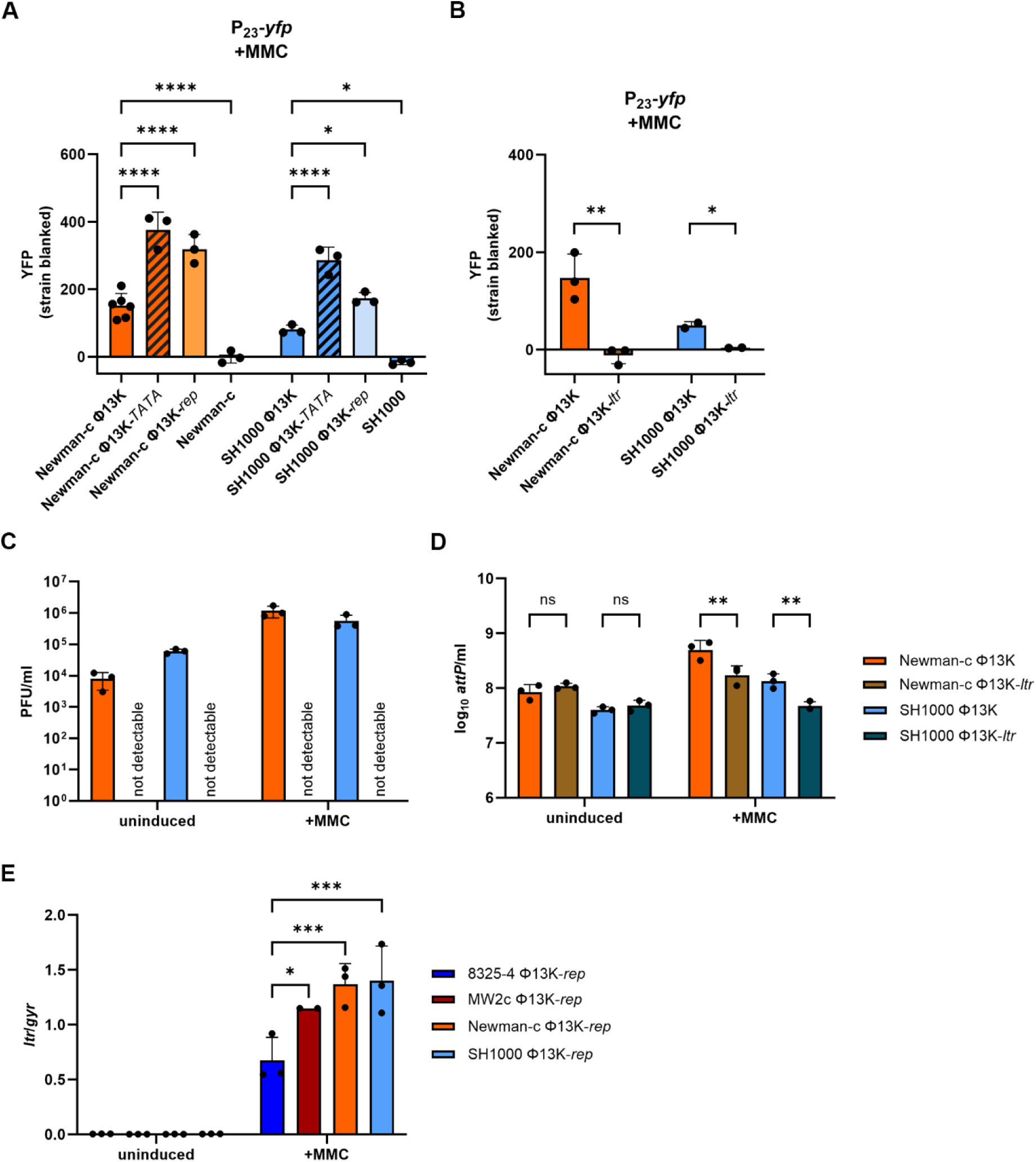
P_23_ promoter region is controlled by the phage-encoded late transcriptional regulator (Ltr). (A) Single timepoint measurement of P_23_ promoter activity in single-lysogens carrying Φ13K (wild type), Φ*13K-TATA* (TATA-box mutation), or Φ*13K-rep* (replication deficient), and in phage-free strains. Strains were grown to exponential phase, induced with a subinhibitory concentration of mitomycin C (MMC) (300 ng ml^-1^) and grown for further 4 hours. Bacteria were harvested to OD_600_ of 2 and resuspended in PBS. Fluorescence was measured. Arbitrary units of fluorescence (YFP) are shown; strain-specific background fluorescence was subtracted. Data shown are mean ± SD (n ≥ 3). Statistical significance was determined by ordinary one-way ANOVA. (B) Single time point measurement of P_23_ promoter activity in single-lysogens carrying Φ13K (wild type), or Φ*13K-ltr* (*ltr* deletion). Data shown are mean ± SD. Statistical significance was determined by unpaired t-tests for comparisons within strains. (C,D) Phage replication in strains carrying Φ13K (wild type) or Φ13K-*ltr* (*ltr* deletion) under uninduced and induced (+MMC) conditions. Single-lysogenic Φ13K strains were induced with subinhibitory MMC in exponential phase and incubated for 60 min. Phage numbers were determined by plaque assay (C) and phage genome numbers by qPCR on the attachment site (*attP*) (D) Data shown are mean ± SD. Statistical significance was determined by 2way ANOVA on log_10_ transformed data. (E) Strain specific *ltr* gene expression. Single-lysogenic strains were grown to exponential phase, induced with a subinhibitory concentration of MMC and grown for further 4 hours. Bacteria were harvested for RNA Isolation and RT-qPCR was performed. *ltr* gene expression was normalized to *gyrB* expression. Data shown are mean ± SD. Statistical significance was determined by 2way ANOVA. (****p-value < 0.0001, ***p-value < 0.001, **p-value < 0.01, *p-value < 0.05, ns > 0.05).

In contrast to the analyzed phage mutants, a phage-free background resulted in promoter inactivation, confirming that a phage-encoded factor is required for P_23_ activity (Figure 3A). This effect was observed for all four tested strains (Supplemental Figure S2).

### SAOUHSC_02200 is a late transcriptional regulator of the Ltr family IV

To identify the phage factor activating the P_23_ promoter, we compared the genomic organization of Φ13 with other phages. Since late phage genes are located downstream of P_23_, we mined the phage genome for the presence of late transcriptional regulators (Ltrs), which activate late phage gene expression. Quiles-Puchalt et al. described four families of Ltrs in addition to the RinA homologs identified in *S. aureus* phage Φ11. These Ltrs are characterized as small basic proteins encoded at the end of the early phage gene cluster and positioned directly upstream of the promoter they regulate (26).

Due to the localization of *SAOUHSC_02200* upstream of P_23_ promoter region, we aligned the protein sequence of SAOUHSC_02200 (UniProt ID: Q2FWT0) with those of representative members of the different Ltr families. Sequence analysis revealed 59% similarity to the ORF34 of phage Φ55 (UniProt ID: Q4ZB49), encoding a member of the Ltr family IV, and 100% identity to hypothetical protein PVL_60 of *S. aureus* phage PVL (UniProt ID: O80099) (Supplemental Figure S3). These alignment results support the hypothesis that *SAOUHSC_02200* encodes a late transcriptional regulator belonging to the Ltr family IV (LtrC-homologs).

### *SAOUHSC_02200*-encoded Ltr regulates P_23_ promoter

To verify the role of the putative Ltr in P_23_ regulation, we generated a genomic *ltr-*deficient mutant by introducing a stop codon into *SAOUHSC_02200* without altering the P_23_ promoter region. In the absence of functional Ltr, no P_23_ activity was detected, proving that P_23_ activation is dependent on Ltr (Figure 3B). Similar to the Φ13K-TATA mutant, the Φ13K-ltr mutant failed to produce intact phage particles. However, phage genome replication appeared unaffected under uninduced conditions. Following MMC induction, a greater number of genome copies were detectable for the wild type phage than for the *ltr* mutant phage, for which no increase in phage genome copies was detectable (Figure 3C,D). This is likely because only the wild type phage propagates and lyses its hosts.

After identifying Ltr (SAOUHSC_02200) as a regulator and activator of promoter P_23_, we analyzed the expression of *ltr* in different single-lysogenic strain backgrounds under uninduced conditions and 4 h post MMC-induction. Replication-deficient phage mutants (Φ13K-*rep*) were used to avoid multi copy effects. As expected, *ltr* expression was undetectable under uninduced conditions. Upon MMC induction, strain-specific differences in *ltr* expression were detected, with the 8325-4 single-lysogen showing significantly lower expression than the other strains (Figure 3E). These differences correlated with differences in SigB activity (see Figure 2A).

In summary, these results confirm Ltr (encoded by *SAOUHSC_02200*) as the activator of promoter P_23_ and demonstrate that a functional Ltr is essential for production of intact mature phage particles. The variation in *ltr* expression in different strain backgrounds indicates that host factors, particularly SigB, contribute to its regulation.

### Repeat motifs in the P_23_ promoter region are essential for activation

Previous studies have shown that Ltrs generally bind to four imperfect repeats within their target promoter regions (26). Sequence analysis of P_23_ promoter region revealed three perfect 18 bp repeats and an additional partial repeat corresponding to the first 9 bp (Figure 4A). Deletion of this repeated sequence in the reporter plasmid abolished P_23_ activity (Figure 4B). This confirmed the importance of this repeated sequence for promoter activation, likely due to impaired binding of Ltr in the absence of the repeat motif.

**Figure 4:**
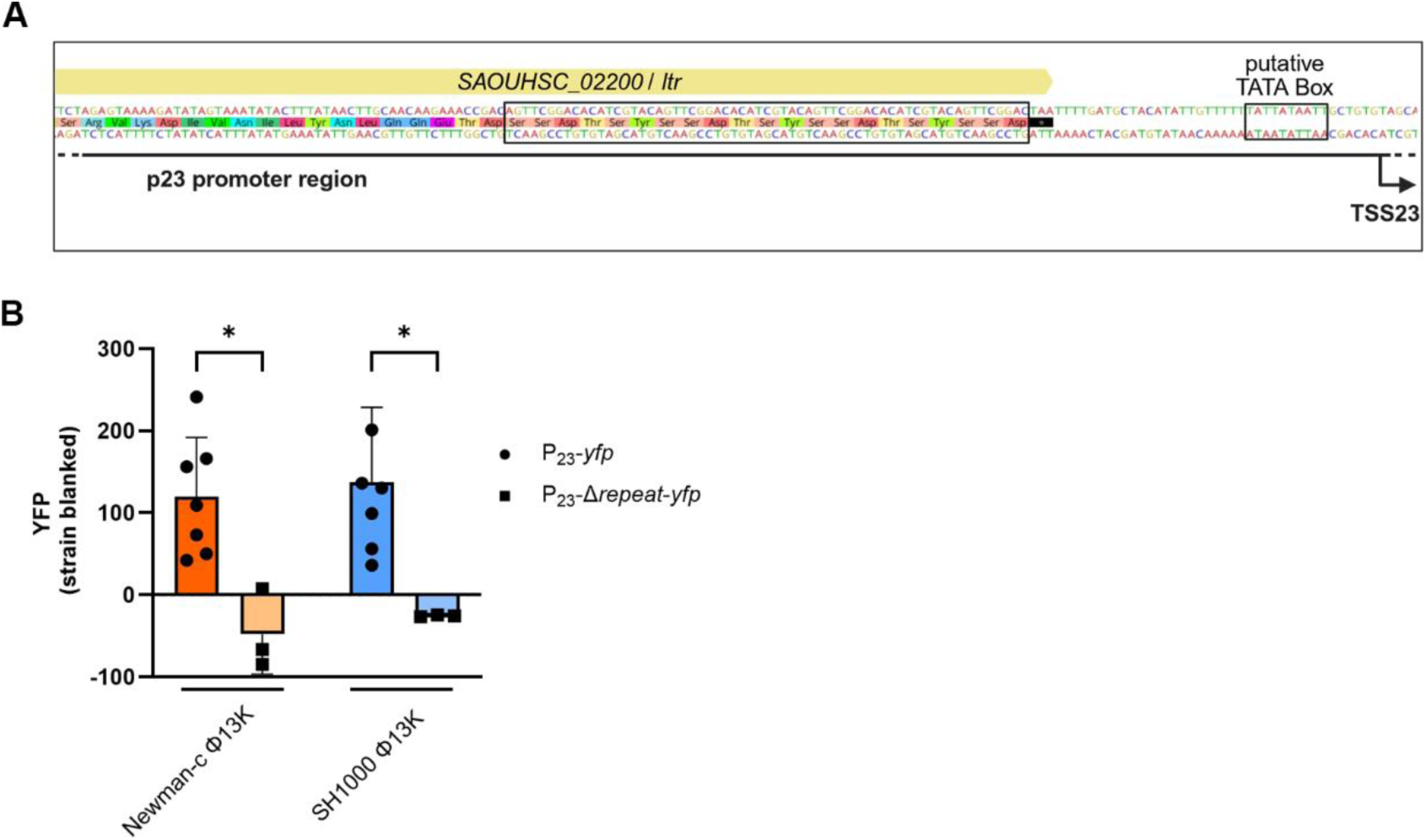
The repeated sequence in the P_23_ promoter region is required for promoter activation. (A) Schematic depiction of repeated sequence in P_23_ promoter region upstream of the putative TATA-box. (B) Single time point measurement of the promoter activity of wild type (P_23_-*yfp)* or mutant P_23_ (P_23_-Δ*repeat*-*yfp*), lacking the repeated region. Single-lysogenic strains were grown to exponential phase, induced with a subinhibitory concentration of MMC and grown for further 4 hours. Bacteria were harvested to OD_600_ of 2 and resuspended in PBS. Fluorescence was measured. Arbitrary units of fluorescence (YFP) are shown; strain-specific background fluorescence was subtracted. Data shown are mean ± SD (n ≥ 3). Statistical significance was determined by ordinary one-way ANOVA. (*p-value < 0.05).

### P_23_ activation by Ltr is essential for late phage gene expression

We next determined the impact of Ltr on P_23_ dependent gene expression. Therefore, gene expression across the phage genome in absence of functional P_23_ or Ltr (Figure 5A) was assessed using RT-qPCR analyses on RNA isolated from single-lysogenic SH1000 strains carrying replication-deficient Φ13K-*rep* or its derivatives with mutated P_23_ (Φ13K-*TATA*), or lacking *ltr* (Φ13K-*ltr*) (Figure 5B-I).

**Figure 5:**
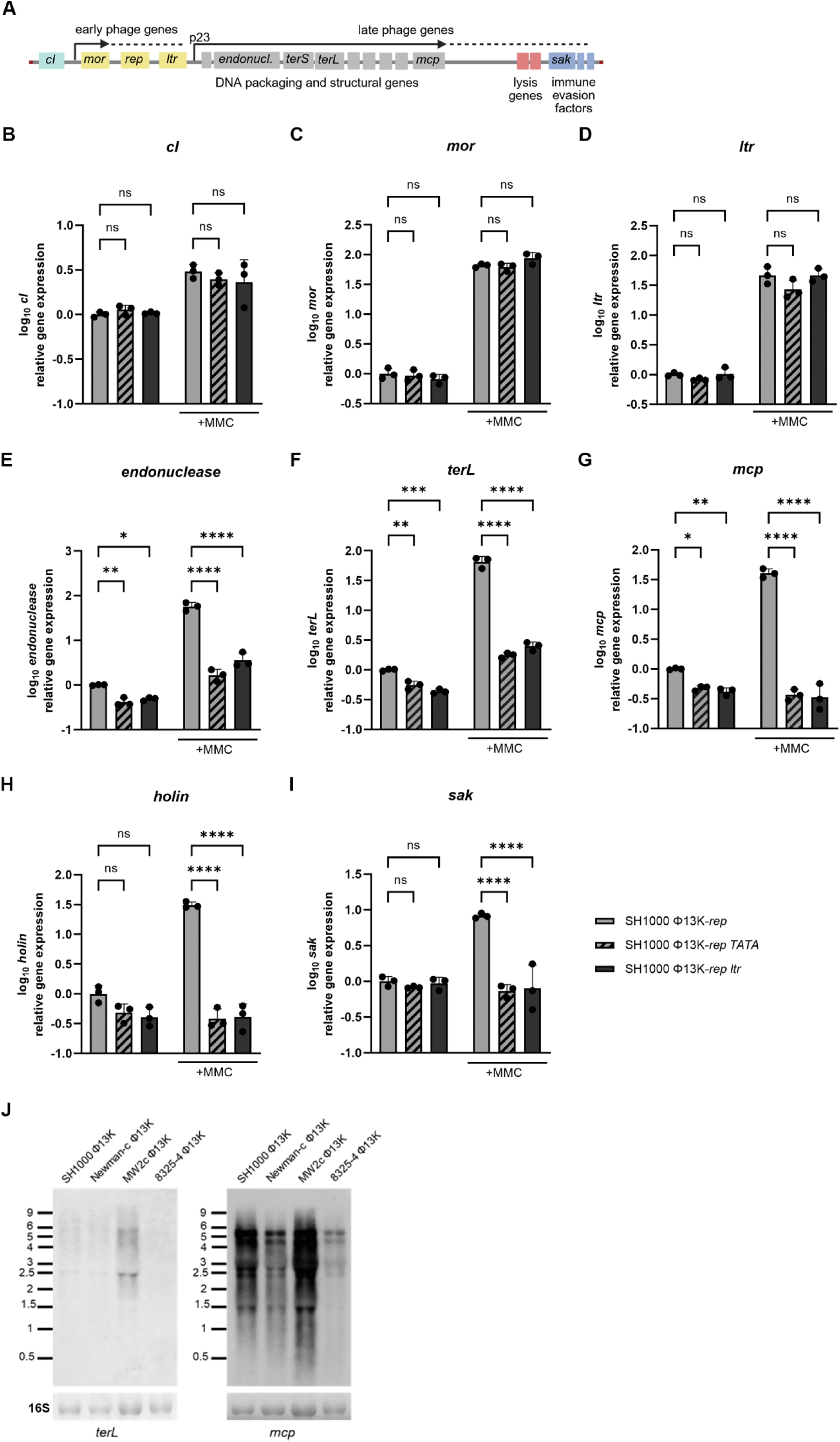
P_23_ promoter activation by Ltr is required for late gene expression. (A) Schematic depiction of the phage genome, indicating early and late phage genes. (B-I) Gene expression analysis of early (*cI*, *mor* and late transcriptional regulator (*ltr*)) and late phage genes (*endonuclease*, large terminase (*terL*), major capsid protein (*mcp*), *holin* and staphylokinase (*sak*)) under uninduced and induced (+MMC) conditions. Strains carrying Φ13K-*rep* (wild type, replication deficient), Φ13K-rep-*TATA* (TATA-box mutation), or Φ13K-rep-*ltr* (*ltr* deletion) were grown to exponential phase, induced with a subinhibitory concentration of mitomycin C (MMC) and grown for 60 min. Bacteria were harvested for RNA isolation, transcripts were quantified by RT-qPCR and normalized to *gyrB* expression by ΔΔCt method.. Data shown are mean ± SD. Statistical significance was determined by ordinary one-way ANOVA on log_10_ transformed data. (****p-value < 0.0001, ***p-value < 0.001, **p-value < 0.01, *p-value < 0.05, ns > 0.05). (J) Northern Blot analysis for *terL* and *mcp* transcripts. Single-lysogens were grown to exponential phase, induced with a subinhibitory concentration of mitomycin C (MMC) and grown for 60 min. Bacteria were harvested for RNA isolation. Blots were hybridized by using digoxigenin-labeled DNA-probes generated by PCR. 16S RNA was detected for loading control. Millenium Marker (ThermoFisher Scientific) was included for size comparison.

Expression of the genetic switch genes *cI* and *mor* were unaffected by disruption of P_23_ or mutation of *ltr*, indicating that early phage gene expression and induction of the lytic life cycle are Ltr-independent. Similarly, *ltr* itself showed no significant change in expression in the phage mutants, suggesting the absence of regulatory feedback on *ltr* expression.

In contrast, the expression of late phage genes involved in DNA-processing and packaging (*endonuclease*, *terL*), as well as the major capsid protein gene (*mcp*), was significantly reduced in both the Φ13K-*rep*-*TATA* and Φ13K-*rep*-*ltr* phage mutants compared to the wild type Φ13K-*rep*. The expression of late phage genes required for bacterial lysis and phage release, such as *holin* was also significantly reduced, alongside the expression of the immune evasion factor staphylokinase (*sak*), located downstream of the lysis genes.

Overall, the expression of all the analyzed late genes downstream of the P_23_ promoter including the immune evasion cluster proved to be dependent on P_23_ and Ltr. Moreover, MMC induction did not induce late gene expression in the mutants, confirming that Ltr is required for late gene activation during the lytic cycle.

Previous studies have shown that late transcriptional regulators activate the transcription of a single large transcript covering all late phage genes (26). Northern Blot analysis using DNA-probes targeting transcripts of *terL* and *mcp* revealed multiple transcripts in different sizes, including a shared transcript of ∼ 6 kb, which indicated co-transcription of *terL* and *mcp* from the P_23_ promoter. However, no larger transcript covering all late phage genes was detected (Figure 5J).

Although additional transcriptional start sites and transcriptional units were identified for late genes (24), our data indicate Ltr as major phage regulator of late phage gene expression from the P_23_ promoter.

## Discussion

The impact of host factors such as RecA or SarA on Φ13 excision and replication have been previously described (22, 27). Here we focused on regulatory mechanisms controlling phage assembly. We identified Ltr as a key regulator of late gene transcription. Ltr, encoded by the final gene on the early phage module, activates P_23_ and thereby initiates the expression of genes involved in capsid assembly, DNA processing and packaging, lysis, and immune evasion. Both *ltr* expression and P_23_ activity exhibit strain-specific differences, which appear to be dependent on the alternative sigma factor SigB and its downstream effector SpoVG. We propose that Ltr could be a regulatory target for the switch between the lytic phage life cycle and active lysogeny (Figure 6).

**Figure 6:**
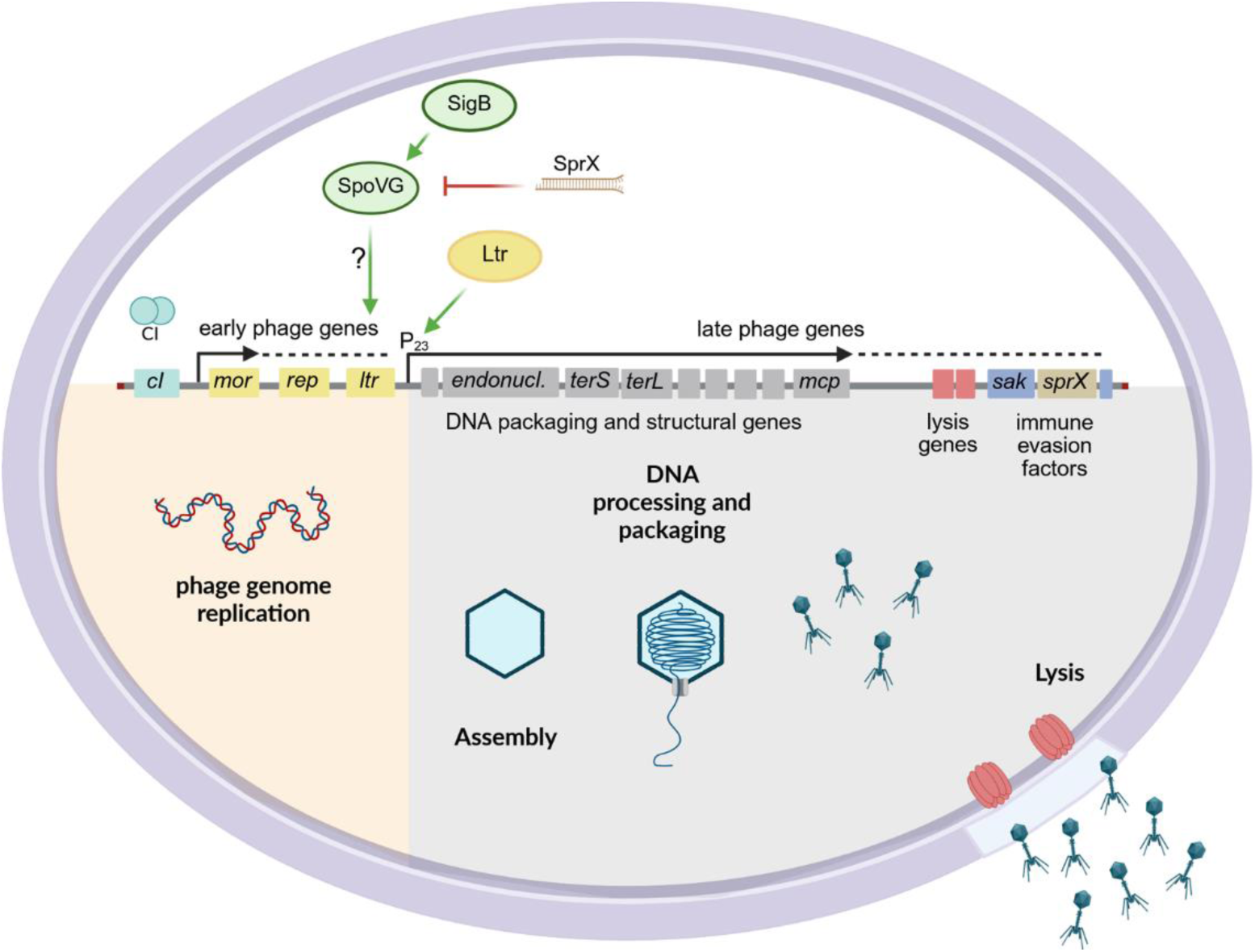
Temporal control of phage gene expression within the bacterial host. Genetic organization of the bacteriophage Φ13. Release of CI from the *mor* promoter results in derepression of the lytic cycle and expression of early lytic genes, including the replication factor (*rep*) and late transcriptional regulator (*ltr*). Subsequently, late expression of DNA-packaging (*endonuclease*, *terS*, *terL*) and structural genes (e.g. *mcp*) leads to DNA-processing, capsid assembly and DNA-packaging. The products of lysis genes enable host cell lysis, releasing mature phage particles, while immune evasion factors (e.g. *sak*) aid in escaping the host immune system. These late phage genes are transcribed from the P_23_ promoter, which is activated by the Ltr transcriptional regulator encoded by *ltr*. Ltr expression and P_23_ activity are regulated in a SigB and SpoVG-dependent manner. Small RNA SprX provides a feedback for post-transcriptional SpoVG repression.

### Ltr (SAOUHSC_02200) regulates late phage gene expression in Φ13

Phage gene regulation is tightly temporally controlled to ensure that genome replication is complete before particle assembly and host lysis occur. The first step in genetic activation is the transition from the lysogenic to the lytic life cycle. In the model phage λ, this genetic switch is regulated by the interplay between the CI repressor and Cro anti-repressor. Similarly, in the Sa3int phage Φ13 this switch involves a cleavable CI-repressor and the small anti-repressor Mor, originally described in lactococcal phages (27, 28). Activation of the SOS-response induces RecA-dependent auto-cleavage of CI, leading to de-repression of early lytic genes and phage genome replication. Subsequently, late phage genes are expressed to enable the assembly of mature phage particles, and bacterial lysis.

In *S. aureus* phage Φ11, the expression of late genes is controlled by the transcriptional regulator RinA, located upstream of the *terS* gene. Deletion of *rinA* abolishes the production of functional phage particles and reduces late gene expression (29). Quiles-Puchalt et al. described four additional families of late transcriptional regulators (Ltrs), which share the size of about 130 aa and a similar localization within the phage genome, although they show no sequence similarity to RinA or known bacterial transcription factors (26).

Here, the Ltr of Φ13 was identified based on the localization upstream of the endonuclease gene and the predicted TSS23. Mutation of *ltr* or P_23_ abolished the production of functional phage particles and significantly reduced the expression of all genes downstream of P_23_. Moreover, P_23_ activity was completely lost in phages lacking a functional *ltr* gene, confirming Ltr as the key regulator of late gene expression in Φ13. Sequence alignment to the different Ltr families revealed 30% identity and 59% similarity with ORF34 of phage Φ55 (Ltr IV family) and 100% identity to hypothetical protein PVL_60 of Staphylococcal phage PVL. Panton-Valentine leucocidin (PVL), the bi-component pore-forming cytotoxin encoded by phage PVL and several other phages, contributes to the virulence of *S. aureus* by targeting and lysing host immune cells such as macrophages and neutrophils (30, 31). Comparative genome analysis revealed extensive sequence identity between Φ13K and staphylococcal phage PVL (Supplemental Figure S3). For instance, one 19 kb region spans from the P_23_ promoter region up to a locus upstream of the lysis genes. Another conserved region includes the genetic switch genes *cI* and *mor*. Phage genomes are known to have interchangeable modules fulfilling the same functions, which lead to mosaicism within phage genomes (3). The high degree of conservation between Φ13K and staphylococcal phage PVL in regions important for transcriptional regulation, suggests a similar phage-host interaction for both phages in *S. aureus*.

Previous studies have shown that repeated sequences in the promoter region serve as recognition targets for Ltr binding (26, 29). In *Lactococcus lactis* phage TP901-1, sharing the anti-repressor MOR with Φ13, the late transcriptional regulator Alt binds to four imperfect direct repeats located -76 to -32 bp upstream of the transcriptional start site (32, 33). The promoter region downstream of Φ13-Ltr also consists of four repeats located at the 3’ end of the regulator’s coding sequence. Deletion of this repeat region, between position -107 and -42 bp relative to the TSS, abolished P_23_ activity, confirming that binding of Φ13-Ltr to this sequence is required for promoter activation.

### Transcriptional organization and regulation of late phage genes in Φ13

For both RinA- and Ltr-homologous regulators, transcription from their respective promoters results in a single transcript for DNA-packaging, structural and lysis genes (26, 29). Consistent with this, Northern blot analysis confirmed co-transcription of *terL* and *mcp*, however, no transcript spanning across the phage genome from P_23_ to the immune evasion cluster was detected. In Φ13, additional transcriptional start sites were identified downstream of TSS23, indicating a more complex regulatory network. These include TSS located upstream of *mcp* (TSS25, 26), lysis genes (TSS30) and immune evasion factors (TSS36) (24). Previous work suggested co-transcription of the lysis gene *amidase* and the immune evasion factor *sak* (34). Transcript size analysis by Northern blot revealed a transcript of 3 kb hybridizing with a *sak*-specific probe, confirming the presence of additional transcriptional start sites within the late genetic modules. The major regulatory factor, however, appears to be *ltr*, as phages lacking *ltr* or carrying disruptions in P_23_ did not express late phage genes, including *sak* and lysis genes. These findings suggest that potential additional regulatory processes are dependent on initial Ltr activation of promoter P_23_.

### Regulatory mechanisms controlling Ltr and P_23_ activity

The post-transcriptional inhibition of LlgA (a Ltr family protein) activity has been shown to block late gene transcription in *Listeria monocytogenes* phage Φ10403S in the intracellular niche. This interruption of the phage replication process induces a pseudo-lysogenic state that allows prophage excision and subsequent *comK* expression required for phagosomal escape (35). Although the bacterial factors mediating LlgA regulation remained unidentified, it was proposed that LlgA activity is under host-dependent control for beneficial regulation of the phage life cycle in different stages of infection.

We observed strain-specific differences in P_23_ activity and *ltr* expression pattern also indicating a strong impact of bacterial host factors on the fate of the phage. Interestingly, SigB activity within the different strains correlated with *ltr* expression differences, indicating a regulatory effect of the host factor SigB on *ltr* expression. Φ13 or PVL phage Sa2mw integrated into the SigB deficient 8325-4 lysogens were previously shown to exhibit reduced mobility compared to other SigB-positive strains (24, 25). RinA, the late transcriptional regulator of Φ11, exhibits the highest expression in the late-exponential phase (36) which is consistent with SigB activity during later growth phases (37).

However, no SigB consensus sequences could be identified in the promoter initiating *ltr* or P_23_. Deletion of *spoVG* resulted in reduced P_23_ activity and phage assembly similar to SigB deletion. SpoVG acts as a downstream effector of SigB and was proposed to activate transcription of SigB-dependent genes lacking a SigB consensus sequence (38, 39). Thus, SpoVG likely mediates the impact of SigB on phage gene regulation. SpoVG is broadly conserved, especially among Gram-positive bacteria. *L. monocytogenes* SpoVG was shown to bind RNA with a greater affinity than DNA and it was suggested that the protein is mainly acting as RNA-binding protein thereby functioning as a global post-transcriptional gene regulator (40).

SpoVG in *S. aureus* is post-transcriptionally suppressed through interference with a small RNA, SprX (41), which is located within several Sa3int phages downstream of *sak*. Thus, there seems to be a strong feedback mechanism whereby upon phage replication the increase of SprX likely restricts SpoVG-dependent phage assembly. Under certain infectious conditions the SOS response and SigB activity are activated, which then may promote SpoVG activity to overcome SprX restriction and enable the phage to escape from the bacterial host. However, additional host factors likely contribute to the regulation of phage late gene expression. TSS prediction showed additional strain-specific TSSs in MW2c and 8325-4. Furthermore, the high P_23_ activity of MW2 compared to other SigB-positive strains could not be linked to higher SigB activity in MW2 and thus remain unclear.

## Material and Methods

### Growth conditions

Strains used in this work are listed in Table S1. If not stated differently, strains were grown in Tryptic Soy Broth (TSB) (Oxoid) at 37°C and 200 rpm. For strains carrying resistance genes, antibiotics (erythromycin (erm) 10 μg ml^-1^, tetracyclin (tet) 3 μg ml^-1^, kanamycin (kan) 50 μg ml^-1^, chloramphenicol (cm) 10 μg ml^-1^) were added in precultures and for selection on agar plates.

### Generation of strains and mutants

Oligonucleotides and plasmids used for generation of mutants are listed in Table S2 and Table S3.

#### Promoter fusion constructs for promoter activity measurements

By Gibson Assembly, promoter regions of genes of interest (P_gene_) were cloned into reporter plasmids upstream of a strong ribosomal binding site (RBS) followed by a gene coding for a yellow fluorescent protein (*yfp*: gpVenus). To generate P_23_-*yfp* (pCG896) 50 bp downstream of the predicted transcriptional start site 23 (TSS23) and 200 bp upstream were amplified by PCR with the primer pair pCG896gibfor/pCG896gibrev (24). The amplified promoter regions were cloned into the reporter plasmid pCG725 digested with SalI and SphI (24, 42). To generate P_23_-*TATA-yfp* (pCG910) the bases of the putative TATA Box of promoter region 23 in pCG896 were replaced with Cs and Gs by site directed mutagenesis (SDM) using the primer pair pCG910SDMfor/pCG910SDMrev. To generate P_23_-Δ*repeat*-*yfp* (pCG943) the repeated sequence of 65 bp was deleted from pCG896 by SDM using the primer pair pCG943SDMfor/pCG943SDMrev. Site directed mutagenesis was performed according to the manufacturers instructions (Q5 Site-Directed Mutagenesis Kit).

Promoter fusion plasmids were verified by sequencing and introduced into the final bacterial strains by transduction via *S. aureus* RN4220 or direct electroporation. Final strains were verified by PCR on the reporter plasmids.

#### Late transcriptional regulator (ltr) mutant (Φ13K-ltr)

Flanking regions were amplified by PCR using primer pairs pCG926insert1gibfor/pCG926insert1gibrev and pCG926insert2gibfor/pCG926insert2gibrev. The overlapping primers pCG926insert1gibrev and pCG926insert2gibfor contained point mutations to introduce a stop codon into *SAOUHSC_02200*. Both fragments were cloned into BamHI digested pIMAY-Z vector and transformed into *E. coli* DC10B. After verification by sequencing, the plasmid was transformed into the final *S. aureus* strains by electroporation. pIMAY-Z mutagenesis for genomic integration was performed as described before (43).

#### Generation of phage Φ13K-rep

Replication deficient phage mutants were generated as described before (24).

#### Generation of mutants (sigB, spoVG)

For the generation of *sigB*::*tet* and *spoVG*::*erm* mutants, recipient strains were transduced with Φ11 phage lysates, generated by using strains included in Table S1. Deletions were verified by PCR on the flanking regions within the resistance cassettes and on the respective gene.

### Promoter activity measurement

#### Time measurement of promoter activity

Promoter activity over time was analyzed in a Tecan Spark plate reader as following. Cultures were inoculated in 96-well plate (Greiner, F-Bottom, clear) to OD_600_ of 0.05, grown for 2 h at 37°C and induced with subinhibitory concentration of mitomycin C (MMC, 300 ng ml^-1^). Optical density (absorbance: 600 nm) and fluorescence (gpVenus: excitation 500 nm, emission 545 nm) were measured in 30 min time-intervals for 24 hours. Fluorescence measurements were normalized to OD_600_ and strain-background fluorescence was subtracted.

#### Single time point measurement of promoter activity

For single time point measurements of promoter activity, measurements were performed as described before (22). In brief, 4 hours after induction with MMC bacteria were harvested, adjusted to OD_600_ of 2 in PBS and fluorescence was measured in the plate reader using the same settings as mentioned for time measurement of promoter activity. For analysis, strain-specific background fluorescence was subtracted.

### Prophage induction and phage replication

Phage replication following induction was analyzed as described before (24). In short, single-lysogenic strains were grown to OD_600_ of 0.7, induced with a subinhibitory concentration of MMC (300 ng ml^-1^) and incubated for further 1 or 4 h. Subsequently, supernatants were harvested by centrifugation and sterile filtrated (0.45 µm). Analysis of phage replication of transposon strains was performed in 24 well plates shaking at 130 rpm. Phage numbers in the supernatant were quantified by quantitative PCR (qPCR) and plaque assays.

### Plaque assay

Phage titer were determined in the supernatants by the agar overlay method. Phage top agar (Casaminoacids 3 g l^-1^, Yeast Extract 3 g l^-1^, NaCl 5.9 g l^-1^, Agar 7.5 g l^-1^) was mixed with *S. aureus* LS1 grown to OD_600_ of 0.1 and poured onto NB_2_ agar plates supplemented with CaCl_2_. Supernatants were diluted in phage buffer (Tris 50 mM, CaCl_2_ 4 mM, MgSO_4_ 1mM, NaCl 5.9 g l^-1^, gelatine 1 g l^-1^) and spotted onto the bacterial lawn. Phage titers were calculated as plaque-forming units (PFU) per ml, after overnight incubation at 37°C.

### qPCR for phage genome quantification

Quantification of phage genomes within sterile filtered culture supernatants was performed as described before (22). In brief, supernatants were treated with proteinase K and diluted 1:10 in nuclease-free water. QuantiFAST SYBR Green PCR Kit (Qiagen) was used for qPCRs, with primers spanning the attachment site (*attP*) on the phage genome.

Oligonucleotides used for qPCR are listed in Table S2.

### RNA isolation and RT-qPCR

For RNA isolation, bacteria were harvested and resuspended in 1 ml TRIzol (Thermo Fisher Scientific). Cells were lysed using a high-speed homogenizer (6500 rpm) and zirconia/silica beads (0.1 mm diameter). RNA was isolated as recommended by the TRIzol manufacturer. Five µg of the isolated RNA were DNase-treated (Roche) and subsequently diluted 1:10 in nuclease-free water for RT-qPCR.

To determine gene expression of target genes, RT-qPCR was performed with QuantiFast SYBR Green RT-PCR Kit (Qiagen) using the Quantstudio3 system (Applied Biosystems) with the recommended settings. For strain comparisons gene expression was normalized to *gyrB* expression. Relative gene expression, comparing wild type to mutants, was calculated using the ΔΔCT method with *gyrB* as the housekeeping gene for normalization. Uninduced wild type bacteria were used as control condition.

Oligonucleotides used for RT-qPCR are listed in Table S2.

### Northern Blot

Northern Blot analysis was done as described before (44). Transcripts on Northern blots were hybridized with digoxigenin-labelled DNA probes generated by PCR (Supplementary Table S2).

## Statistical analysis

Statistical analyses were performed using GraphPad Prism 10.3.1 software. All data represent values from independent biological replicates. Tukey’s multiple comparisons tests were performed post-hoc for ANOVA tests. P-values < 0.05 were considered significant.

Schematic depictions were created in https://BioRender.com.

## Acknowledgements

We thank Libera Lo Presti for editing of the manuscript. We thank Janes Krusche for helpful scientific discussions.

