## Supplementary Information for "The phage Φ13-encoded transcriptional regulator Ltr controls phage assembly in Staphylococcus aureus"

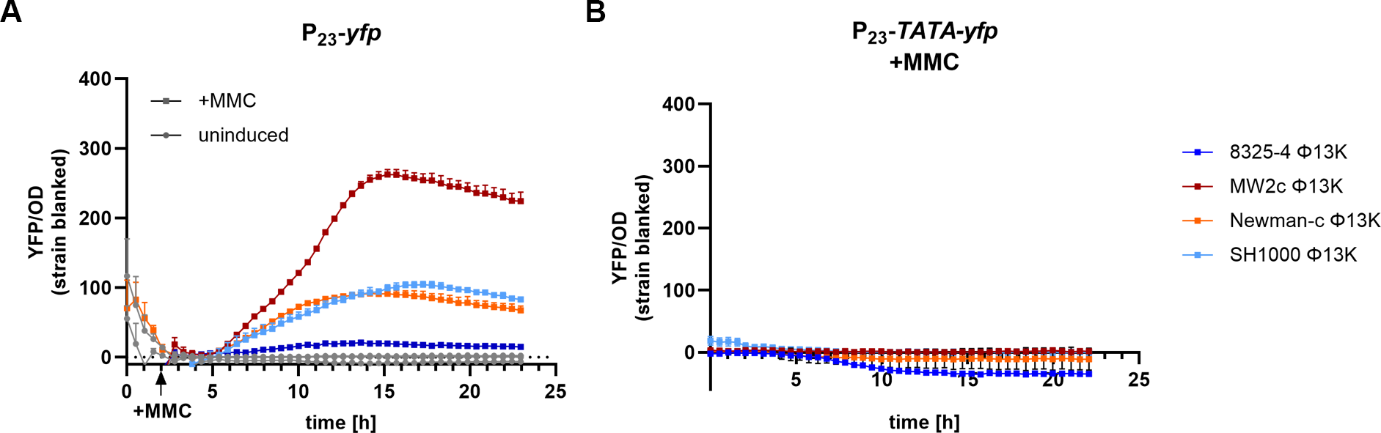

**Figure S1: P_23_ promoter activity measurement over time.** Strains carrying (A) the wild type (P*_23_*-*yfp*: pCG896) or (B) the TATA mutant reporter plasmid (P*_23_*-*TATA*-*yfp*: pCG910) were grown to exponential phase, induced with subinhibitory mitomycin C (MMC), and incubated for 24 h. Optical density and fluorescence were measured. Arbitrary units of fluorescence (YFP) are shown normalized to OD_600_; strain-specific background fluorescence was subtracted. (A) Comparison of strain-specific activity of the wild type P_23_ promoter under uninduced and induced (+MMC) conditions. (B) Promoter activity of P_23_ with mutated TATA-box (P_23_-*TATA*-*yfp*).

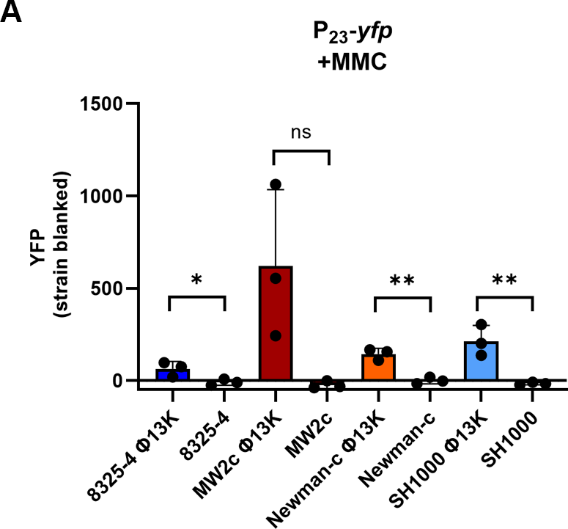

**Figure S2: P23 activity in single-lysogenic and phage-free background.** (A) Single time point measurement of P_23_ promoter activity in single-lysogens carrying Φ13K (wild type) and in phage-free strains. Strains were grown to exponential phase, induced with a subinhibitory concentration of mitomycin C (MMC) (300 ng ml^-1^), and grown for further 4 hours. Bacteria were harvested to OD_600_ of 2 and resuspended in PBS. Fluorescence was measured. Arbitrary units of fluorescence (YFP) are shown, strain-specific background fluorescence was subtracted. Data shown are mean ± SD (n = 3). Statistical significance was determined by unpaired t-tests within strains. (**p-value < 0.01, *p-value < 0.05, ns > 0.05).

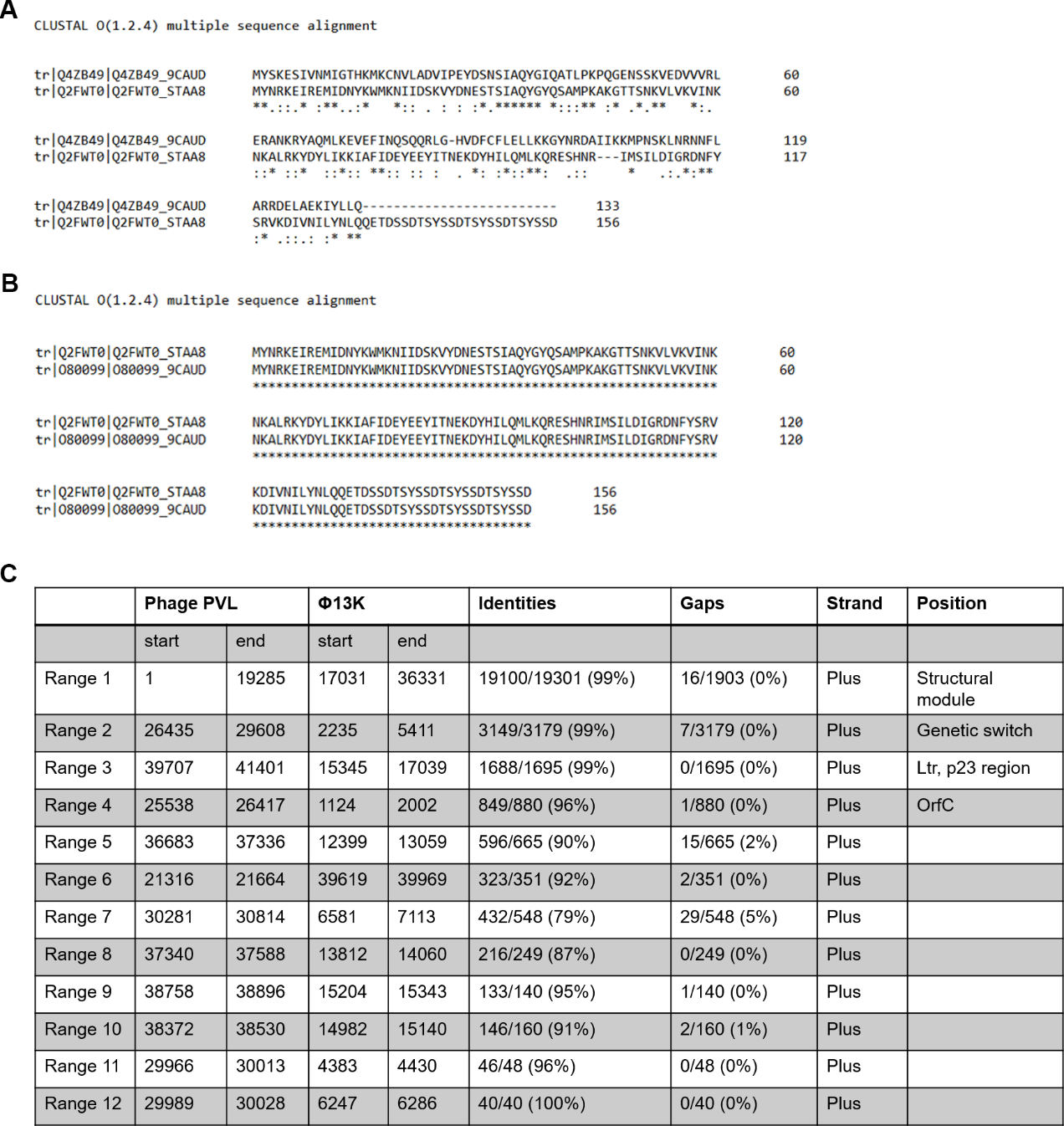

**Figure S3: Alignment of Ltrs and phage genomes** (A,B) Protein sequence alignment of putative Φ13-Ltr (*SAOUHSC_02200*, UniProt ID: Q2FWT0) and ORF34 of phage Φ55 (UniProt ID: Q4ZB49) (A) and hypothetical protein PVL_60 (UniProt ID: O80099) (B). Symbols below sequences indicate the degree of conservation, with ‘*‘ indicating fully conserved residues, ‘:‘ indicating conservation between groups of strongly similar properties (Gonnet PAM 250 > 0.5), ‘.‘ indicating conservation between groups of weakly similar properties (Gonnet PAM 250 ≤ 0.5), and a space indicating non-conserved residues. (C) Overview of nucleotide alignment results of phage Φ13K genome (see Supplement) and phage PVL genome (Accession number: NC_002321).

**Φ13K genome – nucleotide sequence**

>prophage_phi13Kan_TSS_annotated

**CCAGTTTGGATACATAGAAACCTTGTAACAACAGTATTTATTGGGTTTGGAGTCCCTAATGGGTCCCTAAATTACATACTTTCTAAAATTTTAGTTGTTTTTTTGTCCTCTTCATTAAATTTTTCTTCTAACAAATGAGAATACACGGATGTAGTTATTGCTATATTTTTATGACCTAATCTTTTAGAAATGTAATGTATAGATACACCTTTTGCTAGTAAATAAGAACAATGAGTGTGTCTTAATGCGTGCGATGTAATAATTGGTATATTATTGACTCTACAGGCTGATTTCAAAGCATTATTGATAGCCTGAAGGTTAATTATAGATCCGGCTTCTTTGAAAATGTAACCATCATAGCTAATTGCAAATGTACTTATGACGTCCATAATGTGTTTCATATCAGATTTAGCGATACTGATATATCTAGGGGAAGTATCGGTTTTTCGCTCGTCAATAAATATAGTGTTTTTCACTTGGTTGATATGCTCAATCTTTATATTTCTTGCACCACTGACACGACAACCCGTACAAATCATTATGAATAGCGCTAATGATGAACGAGTTCTCTTCTTTCTGACGTGATCTTTTAGTATTTCATATTCAGTTACCGAGATGAATTTTTCTTGTTCTGACTTCGTAGGTTTTCCGGCTTTATAATTAACTTTATAAGCGGGGTTTTTAAAAATAAGTCCATCATATAATGCGTCATCTAAAGCTGACCGAATAGCACCGTTTGTTTTTCTTATAGTTTCTTTTGCGTGTTCTTTTGAATAGTCGTTTATGAATTTCTGATAAACTTGTCTATTTATCTTTGATAACTCCATTTTACCTATTTTATGTTTTTGTATATGTTGTAATGCATTTCTATAATGACGGTAGGTATTTTCTTTAACAACAGGTTGTTTATATGTTTTAATCCAATTTTCGAAGTATTCTGCAAGAGTTATATAGTTATCTATATTAAAACCACTTCTTAACTCATTTAACTTGTCTAGTCCAGCAGAATTAGCTTCACGCTTTGTTCTAAAACCTTTCTTACGGTATCTTTTTCCTTCATGCTTAAATTCATATTGCCATTTTTTACCATCGTAACAACGTGTTTTCATGCGTTCCCTCCTCAAAATTGGCAAAAAATAATAAGGGTAGGCGGGCTACCCAAAATTTAGTACTAGGTACTAAATATGTTATAATAAAATAAAAAGTAGGTGATAAGATGACTCAATTTCTAGGGGCGCTTCTTCTTACAGGAGTTTTAGGTTACATACCATATAAATATCTAACAATGATAGGTTTAGTTAGTGAAAAAAACAAGATTATCAATACTCCTGTATTATTGATTTTTTCTATTGAAACATGTTTGATATGGTTTTATACTTTTATAATTTTTAATAATGTTGATTTAAAAAATTTGAGTTTACTTCAGTTGCTTACAGGTCTAAAAGCAAATATTTGGTTTCTAATTATTTTTGTTTTAACAGTGCTTGTATTTAATCCTTTAATTGTTAAATTCATTATCTGGTTAATTAATGAAACAAGAAAGTTTATGAATTTGGATTGTATAAGCTTATTAGACAAAAGAGACAAGTTGTTTAATAACAACGGTAAACCAGTATTTATAGTTATTAAAGACTTTGAAAACAGAATCATTGAAGAGGGTGAACTTAAAACCTATAATTCAGCTGGTAGCGATTTCGATTTACTAGAGGTTGAGCGACAAGATTTCAAAGTATCTGATTTACCGTCAAACGATGAATTGTATATTAAACATACACTTGTAGACCTTAAACAACAAATTAAATTGGATTTATATTTAATGAATGAATATTAATCTTTTTTCTTAGCTTTTTCTGATAAAGTGCTTTTTAAGTTTTCGCTGGCACCCGGCTTTTCAAAACTTTTGTTTATTGGGTTACTACGAGTAGCTTCTTGTTTTTTGTTTTTATCCGCCATAAAATTCTCACCACCATTCAACGTCTACACTTGTAGGCGTTTTTTGTTTAGTAAAATCATAATGAATCTTCTTTGGTTAACTTATCGCCATCTAATTTTTGTGAAATAAATTCCAAGTATTTACGCGCATTATGTGACGATAAATCTTTAGGTAACTCATAAGTGAATGGTTGATTACCACTAGTTAAAACTTCATATACTATAGTTTCTTTTTTATTTTGCAATTAGTTATTTTCATTATAAACTTCCTTTCAAACACTGCTGAAATAGACGTCTTTTTCAAATAAGCATGATTAATACTTCAATTCTTTAATCCACATATATTTAAAAGTGAGGTAGTAGGTAATAAATATAAGACTTAAAGTTAAGATTGCTTTTTTCATGTCAATTTCTCCTTTGTTTATATTTATATTAAAGCGCTAAATATACGTTATTAATCACAATACAACTTTGCCCATTACTTTAATATCACTAAACGAAGCGACTTTGATATCATCATACTTCGGATTTAGAGATACCAAATTAATATAGTCTTCGCATATATCTACACGCTTGATAAGACTTACTCCATCTAATACAACGAGTGCAATTGTACCATCTTTAATAGAATCTTCTTTCTTAATAAAAGCGTATGTTCCTTGTTTTAACATAGGTTCCATTGAATCACCATTAACTAAAATACAAAAATCAGCATTTGATGGCGTTTCGTCTTCTTTAAAAAATACTTCTTCATGCAATATGTCATCATATAATTCTTCTCCTATGCCAGCACCAGTTGCACCACATGCAATATACGATACTAGTTTAGACTCTTTATATCCATCTATAGAAGTGACTTTATTCTGTTCTTCCAATTGTTCATTTGCATAGTTAAGTACGTTTTCTTGGCGGGGAGGTGTGAGTTGAGAAAATATGTTATTGATTTTTGACATTATCGTTTCATCTTGACGTTCTTCATCAGGAACTCGATAAGAATCTACATCATACCCCATAAGCCACGCTTCACCAACGTTCAGAGTTTTAGAAAGTAGGTAAATTCTATCTTGGTCGGGTGATTGTACGTCGTTAATATATTGAGATAAAGTGCTTTTACTTAAAGATATACCTAGTTTCTTTTGATAAGGTTTCGATTTATTAATGATATCTACTTGTTTTAAGTTTCTTATTTTCATAATGTGTTTAAGTCTATTTGAAACTTTTTCTCTCATTTAGTGCACCTCCGTTTGATAACTTCATAATAAAGCTTGTTGAACAAAAATTCAACAAAAAAGTTCATAAATCATGAATTTTTGTATTGACTTGATTCAAAACAAGGTGTAAAGTATAGTTAAGTTCATGATACGTGAACTTGAGAGGAGGTGCTTTTATGTGTTACGACTACTCACGTTTGAGCGGGAAAATAGTAGAAAAGTATGGCACTCAGTACAATTTTGCAATTGCTATGAAATTGTCAGAGAGAAGTTTATCCTTAAAACTCAACGGTAAAGTTGGTTGGAAAGACAGTGAAATATGGAAAGCTATACAACTACTAGATATACCGGTAGAGAAAATACACTTATATTTTTTTAAAGAAAAAGTTCATGTTATATGAACTTAAGGAGGGGCACAATGGAACAAATCACGTTAACCAAAGAAGAGTGTGTCGAACAATGCATCAATAAAGACTTAAAACTTTTAGATTATCGAGTTCAACAAATTTTAGAAGGTGTTCTATCAGAAAGTACCACATACGGTGATGCAAGAAATAAATTAGAAACATTGAAAATTATTGCTGAATCTCATTTTAAAACCGAACATGCTTCAGTTATTTACAAATTAGCATTGAAAAAGTTAGACGAAAAAATCAACGCCACTCCAATTAAAGAGTGACGGAAAGGGAGGATTTTAAATGTTTAAGGTTTTAAATGATATAAAAACTTCTTTAAAAAACCATCCTTGGGGTTGGAAAGAGCACTTACCTTATTTGCTGATGTTAACTCTGTCACTTGTGGCTCTGATTCTCGGTGTTCTGTCCGCGATTCTATGATAACAGGCTTTATATAGATTCCTTTGTTGGTAGTGACTTTGATAGTCACATCCCATTCCCATATCACTGGATATTCTTCGAGCAAAAAAGTACATTCTACACTTTCATAAGGTCCTAAAGTAAATGGAATGGAGTAGTTTTTATCTTTATATCGTATAGGTTTGAACGTTTTTTGTTCATTTACTTTATTTTTAATATCAAATTCAACGTCAATAACAGAAATGGGAAACTTTGTGAAATTAATAAATGTTATATCGTTGTAACTTGATTTGTCATCGACCAAGTAATTAAAGCTTCTGGTAGGTATAACATCGATGTTAATAGAATCTTTCATATAGTCTAAATAATATTTAAGTGCAGTCAGTAAGAAACTAAAAATTGCGATACAAATCGCGATTATGTCCATACTTATCACCTCCTTAGGTTGATAACAACATTATACACGAAAGGAGCATAAACAATATGCAAGCATTAAAAACAAAATCGAACATCGGCGAAATGTTCAACATACAAGAAAAAGAAAATGGAGAAATCGCAATAAGTGCAAGAGAGTTATATAAAGCTTTGGAAGTTAAAAAGCGTTTTAGCGCTTGGGCAGAAATTAACTTGAAGCATTTCAAAGAAAATAGGGATTTTACAAGTGTACTTACAAGTACGGTTGTTAATAACGGAGCTGTAAGACAACTAGAAGATTATGCTTTAACACTTGATGTAGCTAAACATGTTGCGATGATGTCAGGTACAGAAAAAGGTTTTGATTTTAGAGAGTATTTCATCCAAGTAGAGAAAGCATGGAACAGTCCAGAAATGATTATGAAACGTGCTTTAAAAATTGCTAACAACACAATCAATCAATTAGAAACAAAGATTGAACGTGATAAACCAAAAATTGTATTTGCAGATGCAGTAGCTACTACTAAGACATCAATTTTAGTTGGAGAGTTAGCAAAGATCATTAAACAAAACGGTATAAACATCGGGCAACGCAGATTGTTTGAGTGGTTACGTCAAAACGGATTCCTTATTAAACGCAAGGGTGTGGATTATAACATGCCTACACAGTATTCAATGGAACGTGAGTTATTCGAAATTAAAGAAACATCAATCACACATTCGGACGGTCATACATCAATTAGTAAGACGCCAAAAGTAACAGGCAAAGGACAACAATACTTTGTTAATAAGTTTTTAGGAGAAAAACAAACATCTTAATAGGAGGAACGAACAATGCAAGCTCAAAACAAAAAAGTCATTTATTACTACTATGACGAAGCCGGTAATAGACGACCCGTTAATATTCAATACAACGATGGCTACGACTTAATGATAGACCCGCGTTTTATTGAAATGACGCTTGAAAGACATCCGCATTTAAAAAATAACTTTTATGGATTAATAGATGGAAAAGAATTTAAGTTAGATTAAATTTTTGGAAATGCAAAGGAGGCATAACAAATGTTACAAAAATTTAGAATCGCGAAAGAAAAAAATAAATTAAAACTCAAATTACTAAAGCATGCTAGTTACTGTTTAGAAAGAAGTAACAACCCTGAATTGTTGCGAGCAGTTGCAGAGTTGTTAAAGAAGGTTAACTAAATTAGGCCTTATTATTACTTTTTAGAATGTGAACAATAGGTCGATAAAAAACTTAATAAACAAACTATAGCAACTATCAATGAATTTTGAATATGTAAATCGTTCTCGTTTATATAGTTTGTTACAAAGATTTGAATGTCAGCACCTGCTGCAATGCCATTAGACCATCTTATTAACTTTTTGAAAGGATGTGGAAAATCATTTTCGATACGTTTGACAAATTCATCGTGTCTCTTGTAGGTACTTTGCTCATTTATTGGATAGGTCGAATTGATGGCTTCAGCCAAAGTAGAGATAGCAGTTGGATTGATATAAAAATCTCTAATGGTCTGTTGTGCTTGAAGTACAATCTCATCATCAAACCTATAGAGTTCCTTAAAAGATTTTATCGTTTCTTCAGAAAATAAATTTCTTTGAAATGTTAGAGATGAAAAAGAATTACGCAAATTAAAATTCATTTCAATTAAGTTGTTTAGATGAAAGTCTACTTTGAAGTCAGAAAATAAATTTATGTTGTTTCTATTAATTATATCTAATTGGTACTTAGGTTTTAAAGATTGTTTAATTGCCATACTTTTAGAAATTTCAACATTACTAATTACGTTATTAATAGAAAAACGAACATTTTTTAAAGGATCAATATACACCAATATCACCTCCTTTCACTAGGAGATAACAACATTATACACGAAAGGAAAGATAGAAATGCCACATATTTTAAACGTAACAGTTCCAATACCTGAAACACATGTACTTATCACAAAAGATGAATATGATGAGCTAATTGGTTATTCATTAGACCCTGTATGGAACATGAGTGACTTAAAGAAGAAATTAAAAATTGCATCTGATGAGACTATCAAGGACAGATTACTATTTCATCCTAGATTTGAAAAAGAACTAAGAGCGCAAGGAATTGTGCATTACCCAGATGAGAATTTTAATCGCTGGAGATTTAACGCAAGAAAGATGAATAAATTCGTCGATGAGCATTTCAATGAAATATATAAGGAGAGAATAAAATGAGCAACATTTATAAAAGCTACCTAGTAGCAGTACTGTGCTTTACAGTCTTAGCAATTGTACTTATGCCGTTTCTATACTTCACTACTGCATGGTCGATTGCGGGATTCGCAAGTATCGCAACATTCATATTTTATAAGGAATACTTTTATGAAGAATAAAAAAACTGTTACTCACGGCAATGAGTAACAGTCTAAACAATTAGAAAATTAATGCATATTCAATATAAAACGAAATAAAGGAAGTGTCAACAATGTACTACAAAATTGGCGATGTATGTCAAAAAGTAATTAATGTAGACGGATTCGATTTTAAATTAGCAGTTAAGAAACAAGATTACAGCATTCTAGTGAATGTCTTAGATTTAGAAGATAGATTTATCGACGGTATAAATATAACAGATGAGAATGATCTATACACAGCATTAGACATATTAAATCAATCTATTTATGAATGGATTGAAGAGAACACAGACGAAAGAGACAGGCTAATTAACTTAGTCATGAGATGGTAGGAGGTTGCTATGAAGCAGACTGTAACTTATCTAATCAAGCATAAAGATGAAAATCTATTTATTACAAACCGACCAACTGAAGTGAACGACACAGTGAAGTATTCAACTGATATGCGAGACGCAAGAGAATTCGACGGACTAGACAAAACCGTTATTGATATGTCTAAGCACAAAGCTATTAAGAAAACAGTGACAGAAACAATTGAGTACGAGAAGGTAGAACATGACTGAAAAAACTAATCAAGATGTCGATATCTTAACGCAACTAGGTGTAAAAGACATCAGCAAACAAAATGCAAACAAGTTTTATAAATTTGCGATATACGGCAAGTTCGGGACTGGTAAAACTACGTTTTTAACAAAAGATAACAACGCCTTAGTACTAGATATAAATGAGGACGGAACAACGGTAACAGAAGATGGGGCAGTTGTGCAGATTAAGAATTACAAGCATTTTAGTGCAGTGATTAAGATGTTACCTAAAATTATTGAACAACTCAGAGAAAACGGAAAACAAATTGATGTTGTAGTGATTGAAACAATCCAAAAGCTACGTGATATCACTATGGACGACATCATGGACGGAAAATTAAAGAAACCAACATTTAATGATTGGGGCGAGTGTGCTACACGCATTGTAAGTATTTATCGTTATATTTCTAAATTACAAGAACATTATCAATTCCATCTTGCTATAAGTGGACACGAGGGAATTAACAAAGACAAAGATGATGAGGGTAGCACTATCAATCCAACAATCACGATAGAGGCACAAGATCAAATAAAAAAAGCGGTCATCAGTCAATCTGATGTGTTAGCAAGAATGACAATAGAAGAACATGAGCAAGACGGCGAAAAAGCTTATCAATATGTTCTTAACGCTGAACCATCAAACTTATTCGAGACAAAGATAAGACACTCAAGCAACATTAAAATTAACAACAAACGTTTCATTAATCCAAGTATTAACGACGTAGTACAAGCAATCAGAAATGGAAACTAATAAAAAAACTAAAAAGGACGGTATTTAATTATGAAAATCACAGGACAAGCGCAATTTACTAAAGAAACAAATCAAGAAAAGTTTTATAACGGCTCAGCAGGGTTTCAAGCTGGAGAATTCACAGTGAAAGTTAAAAATATTGAATTCAATGATAGAGAAAATAGATATTTCACAATCGTATTTGAAAATGATGAAGGCAAACAATATAAACATAATCAATTTGTACCGCCGTATAAATATGATTTCCAAGAAAAACAATTGATTGAATTAGTTACTCGATTAGGTATTAAGTTAAATCTTCCTAGCTTAGATTTTGATACCAATGATCTTATTGGTAAGTTTTGTCACTTGGTATTGAAATGGAAATTCAATGAAGATGAAGGTAAGTATTTTACGGATTTTTCATTTATTAAACCTTACAAAAAGGGCGATGATGTTGTTAACAAACCTATTCCGAAGACAGATAAGCAAAAAGCTGAAGAAAATAACGGGGCACAACAACAAACATCAATGTCTCAACAAAGCAATCCATTTGAAAGCAGTGGCCAATTTGGATATGACGACCAAGATTTAGCGTTTTAAGGTGTGGTTTAAATGCAATACATTACAAGATACCAGAAAGACAATGACGGCACTTATTCCGTCGTTGCTACTGGTGTTGAACTTGAACAAAGTCACATTGACTTACTAGAAAACGGATATCCACTAAAAGCAGAAGTAGAGGTTCCGGATAATAAAAAACTATCTATAGAACAACGCAAAAAAATATTCGCAATGTGTAGAGATATAGAACTTCACTGGGGAGAACCGGTGGAATCAATTAGAAAATTATTACAAACAGAATTGGAAATTATGAAAGGTTATGAAGAAATCAGTCTGCGCGACTGTTCTATGAAAGTTGCAAGGGAGTTAATAGAACTGATTATAGCGTTTATGTTTCATCATCAAATACCTATGAGCATAGAAACAAGCAAGTTGTTAAGTGAAGATAAAGCACTATTGTATTGGGCTACAATCAACCGCAACTGTGTAATTTGTGGAAAGCCTCACGCAGACCTAGCGCATTATGAAGCAGTCGGCAGAGGAATGAACAGAAACAAAATGAATCACTACAACAAACATGTATTAGCGTTATGTCGCGAACACCATAACCAGCAACATGCGATTGGCGTTAAGTCGTTTGATGATAAATATCACTTGCATGACTCGTGGATAAAAGTTGATGAGAGGCTCAATAAAATGCTGAAAGGAGAGAAAAAGGAATGAATAGACTAAGAATAATAAAAATAGCACTCCTAATCGTCATCTTGGCGGAAGAGATTAGAAGCGCTAAAAAAATTAAAAAATTTACCCCTGAGGATTCTAAAGGTTTTCCTGATATAACAAAAGATTCAATAAAAGAACCTAAATAAAAATATTATGGTTGATAAAATCCCATTGTTCTTTTGTTAACCACCCTTGTTTGTTATTGACTATTTCTGTAACAAACAGCTTATCTCCAGAATCGAGATAAGGTTTCAACTTTTCTATCATTTCTGAAGTTGATAAAGAAGAACGGAATAAAAATGAAGATTTCCAATAATTGCAATGACCATTAGAAATTTCCTTTTTTATAACATTTCTCAATTCCTCATATTTTTGTCCGGGTGAGTTTAAATCATATGTTAACATATAAGGTTTTTCCATATTTTATTCACCCCCAATCTAACGCAGTAGCGATAACAAAATTATACCAGAAAGGAGATAACGAAATGGCAACATTTAGAGTTTACAAAGAATCAGGTAACTTTGTCACAGTACACAAAGATTTTATACATGATTCTAATATAAGTTGGAAGGCTAAAGGTATTCTACTTTATTTGTTAAGTCGACCTGATAACTGGCAAATTTACGAAACAGAACTAGAGCAACATTCAACTGATGGACTTAGCGGTTTAAAGAGTGGAATCAAGGAACTGGAAGAAATTGGATACATTCAACGTAGTAGAAAACGTGATAAAAGTGGTAGGTTAAATGGTTATGAGTACTTAGTATATGAGCAACCGCACCACATTCGATTTTCCAACGTTGGAAAAACCGTTAACGGTAAAACCAACAATGGAAAAACCGTTAATGGTAAATCGCATACTACTAATAATAATAGTACTAATAATGATTTAACTAATAATAACAATACTAATAATGAAGGAAGTATATTGTCGGGCAACCCGACGGTGTCTTCCATTCCCTATAAAGAAATTATCGAATACTTAAATAAAAAAGCAGGAAAGCATTTTAAACATAATACAGCTAAAACAAAAGATTTTATTAAAGCAAGATGGAATCAAGATTTTAGGTTGGAGGATTTTAAAAAGGTGATTGATATCAAAACAGCTGAATGGTTAAACACGGATAGCGATAAATACCTTAGACCAGAAACACTTTTTGGCAGTAAATTTGAGGGGTACCTCAATCAAAAAATACAACCAACTGGCACGGATCAATTGGAACGCATGAAGTACGACGAAAGTTATTGGGATTAGGGGGATATTATGAAACCACTATTCAGCGAAAAGATAAACGAAAGCTTGAAAAAATATCAACCTACTCATGTCGAAAAAGGATTGAAATGTGAGAGATGTGGAAGTGAATACGACTTATATAAGTTTGCTCCTACTAAAAAACACCCGAATGGTTACGAGTATAAAGACGGTTGCAAATGTGAAATCTATGAGGAATATAAGCGAAACAAGCAACGGAAGATAAACAACATATTCAATCAATCAAACGTTAATCCGTCTTTAAGAGATGCAACAGTCAAAAACTACAAGCCACAAAATGAAAAACAAGTACACGCTAAACAAACAGCAATAGAGTACGTACAAGGCTTCTCTACAAAAGAACCAAAATCATTAATATTGCAAGGTTCATACGGAACTGGTAAAAGCCACCTAGCATACGCTATCGCAAAAGCAGTCAAAGCTAAAGGGCATACGGTTGCTTTTATGCACATACCAATGTTGATGGATCGTATCAAAGCGACATACAACAAAAATGCAGTAGAGACTACAGACGAGTTAGTCAGATTGTTAAGCGATATTGATTTACTTGTACTAGATGATATGGGTGTAGAGAACACAGAACATACTTTAAACAAACTTTTCAGCATTGTTGATAACAGAGTAGGTAAAAACAACATCTTTACAACTAACTTTAGTGATAAAGAACTAAATCAAAATATGAACTGGCAACGTATCAATTCAAGAATGAAACACAATGCAAGAAAAGTAAGAGTAATCGGAGACGATTTCAGGGAGCGAGACGCATGGTAACCAAAGAATTTTTGAAAATTAAACTTGAGTGTTCAGATATGTACGCTCAGAAACTCATAGACGAGGCACAGGGCGATGAAAATAAGTTATATGACCTATTTATCCAAAAACTTGCAGAACGTCACACACGCCCCGCTGTCGTCGAATATTAAGGAGTGTTAAAAATGCCGAAAGAAAAATATTACTTATACCGAGAAGATGGCACGGAAGATATTAAGGTCATCAAGTATAAAGACAACGTAAATGAAGTTTATTCTCTCACAGGAGCCCATTTCAGCGACGAAAAGAAAATCATGACTGATAGAGACCTAAAACGATTCAAAGGCGCTCACGGGCTTCTATATGAGCAAGAGCTAGGATTACAAGCAACGATATTTGATATTTAGAGGTGGCACAATGAGTAAATACAACGCTAAGAAAGTTGAGTACAAAGGAATTGTATTTGATAGCAAAGTAGAGTGCGAATATTACCAATATTTAGAAAGTAATATGAATGGCACTAACTATGATCGTATCGAACTACAACCTAAATTCGAACTACAACCTAAATTTGGGAAGCAAAGACCGATTACGTATATAGCCGATTTCTCTTTGTGGAAGGAAGGGAAACTGGTTGAAGTTATAGACGTTAAAGGTAAGGCGACTGAAGTTGCCAACATCAAAGCGAAGATATTCAGATATCAGTATAGAGATGTGAATTTAACGTGGATATGTAAAGCGCCTAAATACACAGGTCAAGAATGGATGGTATATGAGGACTTAGTGAAAGTCAGACGTAAAAGAAAAAGAGAAATGAAGTGATTTAATGCAACAACAAGCATATATAAATGCAACGATTGATATAAGAATACCTACAGAAGTTGAATATCATCATTTCGATGATGTGGATGATGAAAAAGATATGCTAGCAAAGCGCTTAGATGACAATCCGGATGAATTACTAAAGTATGACAACATAACAATAAGACATGCATATATAGAGGTGGAATAAATGGCGAAAGCAGCAAGAATTGTAAGGATACACGATAAACCTTATAGGTTCAGTAAATTTGAAATGGAATTAATAGAAAGTCACGGTATAACCGCTGGAATGGTTTCTAAGAGAGTAAAAGACGGTTGGGAACTACATGAAGCAATGGACGCACCAGAAGGTACGCGTTTAAGCGAGTACAGAGAAAAGAAAACAATAGAAAGACTGGAACAAGCTAGACTCGAACGCAAATTGGAAAGAAAGCGAAAGAGAGAGGCTGAGCTAAGAAGAAAGAAGCCACACTTGTTTAATGTACCTCAGAAACATTCACGTGATCCGTACTGGTTTGATAATACTTATAACCAAATGTTCAAGAAGTGGCAGGAAGTATAAATGCCTAAAACCGATAGCGCATGTAAAGAATACTTAAACCAATTTTTCGGCTCTAAGAGATATTTGTATCAGGATAACGAACGAGTGGCACATATCCATGTAGTGAATGGCACTTATTACTTTCACGGGCATATCGTACCAGGCTGGCAAGGCGTGAAAAAGACATTTGATACAACCGAAGAGCTCGAAACATATATAAAGCAACATGGTTTGGAATACGAGGAACAGAAGCAACTAACTTTATTTTAAGGAGATGGAAATAATGAAAATCAAAACTGCAAGCATAGAGGTCGAAAAAGTGGAGGTAGTAGTATGATGCCGAAATTTAGAGCGTGGGATAAAGATAAAAAAGTTATGAGTTTTATTGACGAAATCGATTTTAATAGTGGGTACATTTTGATTTCAACAGGTTATAAAAGTTTCAATGAAGTAAAACTATTACAATACACAGGATTTAAAGATGTGCACGGTGTGGAGATTTATGAGGGGGATATTGTTCAAGATTCTTATTCCGGAGAAGTAAGTTTTATCGAGTTTAAAGAAGGAGCCTTTTATATAACTTTTAGCAATGTAACTGAATTAATAAGTGAAAATGACGATATTATTGAAATTATTGGAAATATTTTTGAAAATGAGGAGCTATTGGAGGTTATGAGATGACGGTCACCTTATCAGATGAACAATATAAAAACCTTTGTACTAAATTAAACAAGTTATTAGGTAAATTTCACAAAGCATTAAAAGAACGTGATGAGTACAAGAAGCAACAAGATGAGCTTATCGTGGATATAGGTAAGTTAAGAGAACGTAACAAAGAGTTGGAGAACATGTGGCGCACTCTTAAAAATGAATTGCTTGGAAGATACGAACATTACTGTTTTAAATTTAGAGAACTACACCCTGAGAGCAAAGCGAACAGGATAGGAGCTCTCTATATAGGAGGTAAAAGCACTGCAGATATTATAATGTCGCGAATGGAAGAACTAGACGGAACAAATGAGTTCTACGAATTTTTAGGGCAAATGGAGGAAGACACAAATGAATAACCGTGAACAAATAGAACAATCCGTTATAAGTGCTAGTGCGTATAACGGCAATGACACAGAGGGATTACTAAAAGAGATTGAGGACGTATATAAGAAAGCGCAAGCGTTTGATGAAATACTTGAGGGAATGACAAATGCTATTCAACATTCAGTTAAAGAAGGTATTGAACTTGATGAAGCAATAGGGATTATGGTAAGTCAAGTTATCTATGAATACAAGGAGGAACTGGAGAATGAAAAAATTTAATGTTCAAATCACATATACAGGCATGATTGAAGAGGCTATCGAGGCTGAAAGTTTAGAAGAAGCAGAATTTGAGGCTCATGATATTGCGAGAATGGAAGTGCCATTTGATTGTGATGAATTTGAAATTAATGTAGAGGTGGAACAGGAAAATGAATAACACATTAACAATTGATCAATTACAAGAGTTATTACAAATACAAAAAGAGTTCGACGATAGAATACCGACGCTGAACTTACGAGATAGCAAGATTGCATATGTAGTTGAATTCTTTGAATGGTTTAATACATTGGAAACGTTTAAGAACTGGAAGAAGAAACCAGGTAAGCCGTTAGACGTACAACTTGATGAATTAGCTGACATGTTGGCGTTTGGGTTGAGTATTGCGAATCAAGTAGGAGTGTCATCAGAAGAGATAAAAGAAGCGATTGAATCAAGTTTTAAAAATACAGAATTTCACAAAATGTTTAATTTTAAAGATAAAGAATTTGCTCAAGACGCAGTTGTTAGTACACCACAGATAATATTCAAAGAATTTTATCCCGACCAATTGGCAATTGTAATAGTGATAGACATAGCTTACAACTTATATTCTATCGACCAACTCATTGACGCATACAAAAAGAAAATGAAAAGGAACCACGAAAGACAAGATGGAACAGCAGACGCAGGAAAAGGATACGTGTAAAGACATCTTAGATCGAGTTAAGGAGGTTTTGGGGAAGTGAGAGAACGCACTAAAATTATATATCGTGGTTGGAACAAGGAGATATTTATTTTACAGGGTAAAAATATGAATGTTATTGGTTTGCGCCAAATATTTGATGAACTCAAAAGATTGTACGAAGGTTATAAAATCGTTGTTATTCCAATAGAAGTTGATTTTGAAATCAAATAAATAGGAGTGATGAGAAGTGACACAATACTTAGTCACAACATTCAAAGATTCATCAGGACTACCACATGAACATTTTACTGCTGCTAGAGATAATCAGACGTTTACAGTTGTTGAGGCGGAGAGTAAAGAAGAAGCGAAAGAGAAGTACGAGGCACAAGTTAAAAGGGATGCAGTTATTAAATTAGGTCAGTTGTTTGAAAATATAAGGGAGTGTGGGAAATGATTAAGCAAATATTAAGATTATTATTCTTACTAGCGATGTATGAGCTAGGTAAGTATGTAACTGAGCAAGTATATATTATGATGACAGCTAATGATGATGTAGAGGCGCCGAGTGATTACGTCTTTCGAGCGGAGGTAAGTGAGTGATGTGGATTACTATGACTATTGTATTTGCTATATTGCTATTAGTTTGTATCAGTATTAATAGTGATCGTGCAAGAGAGATACAAGCACTCAGATATATGAATGATTATCTACTTGATGAAGTAGTTAAAACTAAAGGATACAACGGGTTAGAAGAATACAGGATTGAATTGAAGCGAATAAATAACGATATTAAAAAGTAATTTATATTATCGGAGGTATTGCATGTATAACAGGAAAGAAATACGTGAAATGATAGATAACTACAAGTGGATGAAGAACATAATAGACAGTAAAGTCTACGATAACGAAAGTACATCAATTGCACAATATGGTTATCAATCTGCGATGCCAAAAGCTAAAGGCACGACTAGCAATAAAGTGTTAGTGAAAGTTATAAACAAAAACAAAGCGCTTAGAAAGTACGATTACTTGATTAAGAAGATAGCGTTCATTGATGAATATGAAGAATACATCACGAATGAAAAAGATTATCATATTTTACAAATGTTAAAACAACGAGAAAGCCATAATAGGATCATGAGCATTCTTGATATAGGCAGAGACAATTTTTATTCTAGAGTAAAAGATATAGTAAATATACTTTATAACTTGCAACAAGAAACCGACAGTTCGGACACATCGTACAGTTCGGACACATCGTACAGTTCGGACACATCGTACAGTTCGGACTAATTTTGATGCTACATATTGTTTTTTATTATAATTGCTGTGTAGCAAAACATTTATATTTCTTTTGAACTCTCACATTAAGTGAGGGTTTTTATTTTTATAAACAAGAGGTGGAGAATGGAGATATCAAAGTACCAAGAGATAGCTACACGTACACACAATGATGAATTGAATTTAAATGAATATATTACTTGTTACGGCTTAGGTTTAACTCAATCTACAGGCAATGTTACAGATCTAATTAAACAGCATATGTTTTGTAATGTACCGATAGATAAAGGAATTATGATAAATGAACTTAGCGAAGCATTGTGGAATATAGCTAATCTTACTAACGTGTTAGGTATTAACTTGGATGAGATAGCTGGTCATAGTGTTAACACTATCTTGATGAATAAACCTAATCAGACTATCAATTTAGACAATGGTATAAAACGAGGAGACAAAGTATTGTTTCAAGGTAGTAAGTATCTTGTTGATGGATCGATAGGAAACTTATTGTTAATTAGCAATGATAAAGATGATAGACAAGTAACTGTGCAAGATGTTAAGAAAGTCGACAAGGAGTGATGTGCATTGTCTATTATGAAGCGATGTGGTCATCCAACATGTAATGTATTGATTAATCATAATGAAAGTTATTGTGATAAACACAAGCAATATGCAAATGAAAATTACAATGATTTGAGACGTCGAAACGATCCAGAGTATTTAAGATTTTATAAATCGAAAACGTGGCAAAACATGCGTCGAATTGTATTGTTAGAACATGATTTTATTTGTGTTTCTTGTGGCAATCAAGCGACTATGGTTGACCATATTGTACCAACAAAAATTGATTGGGCAAGAAGATTAGACAAAAGTAATTTACAGCCTTTGTGTGATGCTTGCCATAACCAAAAGACAAAAGAAGATTTGAAGAAATATTAAAAAAGATAAAAATAGGAAGTCACCCCAAAGATGAAACGGGCGTCAATGAAAGGTTCTGGAGAACGGAGCAGAGTTTTCTTCTCAAAAAATTCCCTTTATTTAAGTTTTTTTAGTAGGAGGTGCTAATTTATGGCGGGTAGACCTAAGAAGCTTTTGTCAAATTCGAACAAGAATTATACAAAAGAAGAAATTATTGAAAAAGAGCGTCAAGAAGCTCAATTAAATAAATTTTCTAAAATCGATACTGAACCACCGCACTTTTTAGATGAAATAGCGAAACAAGAATACTTAAGAATATTACCGCACATGCAAGAATTGCCAATTTCCAACTTAGATAAAGCACAATTAGCACAATATTGTAGTTTTTATAGTGACTTTGTTAAAGCAAGTTTGATTTTAGAGCGCGAAGACTTGATTTTAGAAGACGACAAAGGAAATCAAAAGGTTAATCCGGCGTTCAACATAAAGGAAAAAGCGGGTATTCGATTGCAACAAACAGCTAATACTTTAGGATTAACTATTGATAGCCGATTGCGTATTATGGTTCCTGATGAAAAAGAAGATGATGATCCATATATGGAATTTGTGAGTGATTAGTAATGACTGATTATGTTACTAAATACGCAAAAAAGGTAGTTTCAGGAGAAATTTTGGCAAGTTTGAAGAATATTCAAGTATGTAAACGTCACCTATCTTTTATGGAGAACCCGCCGAATGGTTGCCATTGGGATAATCATTTGTCTAACAAAGCAATTAAATTTGTGGAAATGCTTCCAGACCCTAAAACAAACCAGCCCATGCCTCTTATGGAGTTTCAGAAATTCATTGTTGGGAGCTTATACGGCTGGCGTAGAGGTCAATACAGAATGTTTACTAAAGCTTATATAAGTATGGCTAGAAAACAAGGTAAGTCTCTAATCGTATCGGGAATGTCCGTTAACGAACTGTTGTTTGGACAATACCCTAAATTTAATAGACAAATTTATGTAGCTTCATCTACTTATAAGCAAGCGCAAACAATATTCAAGATGGCAAGCCAACAAGTAAACCTAATGCGAAGTAAAAGCAAGTTTATCCGTGAAAAAACAGACGTAAGAAAGACAGACATTGAAGATGTATTAAGTAGTTCAGTGTTTGCACCTCTTTCCAATAACCCAGATGCGGTTGATGGTAAAGATCCTACAGTTGCTATTTTGGACGAATTGGCAAGTATGCCTGATGATGAGATGTACTCAAGGTTTAAAACAGGTATGACATTACAAAAAAATCCTTTAACCCTACTTGTTTCAACGGCCGGAGACAATTTAAATAGTCAAATGTACCAAGAGTATAAGTATATTAAACGTATTTTAAATGAAGAAGTAAGAGCTGATAATTACTTTGTATATTGTGCTGAAATGGATTCACAAGAAGAAGTTCAAGATGAAACAAAGTGGATTAAAGCAATGCCGCTTTTAGAATCAAAAGAACATAGAAAAACTATACTTCAAAATGTAAAAGCTGATATACAAGACGAATTAGAAAAAGGGACATCATATCATAAGATTTTGATTAAAAACTTCAATTTATGGCAAGCGCAAAGAGAAGATAGCTTGCTAGATATTTCAGATTGGGAACAAGTAATAACGCCTATGCCTAATATCAATGGTAAAGATGTGTATATAGGTGTCGACTTATCGAGATTGGATGACTTAACATCTGTAGGGTTTATTTTCCCTAACGACGATAAAAAAGTGTTTTTACATAGTCATTCTTTCATTGGATTAAGAACAAACTTAGAACAAAAATCTAAGAGAGACAAAATAAATTATGAATTAGCGATTGAACGTGGCGAAGCTGAGACTACACAATCAGATAGCGGCATGATTGATTATAAACAAGTTATCGATTTTATAGTGAAATTTATAACGACGCATGACCTGAATGTACAGGCTGTTTGCTATGACCCTTGGAATGCGCAAAGTTTTATAACAACAATCGAATCAATGGCTTTAGATTGGCCACTCATTGAAGTGGGACAAAGTTTTAAGGCGTTATCACAATCTATTAAAGAATTTAGAATGTGGGTTGCAGATGAAAGAATACAGCATAACGATAATATGTTACTTACAACATCAGTTAATAATGCCGTTTTGATTCGTGACGGAGAAGACAATGTGAAAATAAATAAAAAAATGAATCGTCAAAAAATAGATCCGATTATTTCGATTATCACAGCTTTCACTGAAGCTAGAATGCACGAATTCCAAGAAAATTGGACGGAGAAATATGAAAGCGAAGAATTCGGATTTTAAAGGTGGTGACAAAATGGACTTGAATAAAATAAATGTCTTTTTTAATTTCTTGGTTGCTAATTTGGTTAGCATCCTTTTTTTATTAGGTTTGTTTGTGGTTAATGTTTCTGTGTATAAAGCATTCGGTCAAAATATAGGACTTTTATGCATTGGTATAACACTGATTGTTATTTCGTTGATTTTAAATCACGAAAGCAATCAAGAAAGGAGTTAGTAGTTGTGGGGATTTTTTATAAAAATGAAAAACGAGACTTGCAATACAACGAAGATGATTTGCAAATGATGGTTCAAACTTTGCCAGGTTTTCAAGGAACAAAATTACGACAATATAAAGATATAGAAGCAATTAGGCATAGCGACATCTTTACGGCAGTTATGATGATTGCTTCTGATTTGGCGCGCATGCCAATTAGGGTGACAGTGAACGGCCAAATTAATTATAGTGACAGGATTGTTAATTTGTTAAATACACGTCCTAACCCAATGTATAACGGCTATATATTCAAATTAGTAGTGTTTGTTAGTGCCTTACTAACATCGCACGGCTATATTGAAATTACACGTGATAAAACAGGAGAACCTATGAATTTAACGTTCAGAAAGACATCCGAAATAGAATTGAAATCAGACGCAAGAGGTCGACTGTATTATTTTCATCAAAGGATAGACAGTAACGGAAATAATATAGAACGTAATGTTAAGTTTGAGGATATGCTAGACATCAAATTTTATTCGTTGGATGGTATAAATGGTTTGTCACTGTTAGACACATTAAGTCGCACGATAGAATCAGATAACAATGGAAAAGATTTCCTTAATAATTTCTTGCGAAATGGCACACATGCTGGTGGTATTTTGAAAATGAAAGGTGTATTAGATAATAAAAAAGCAAGAGACCGTGCCAGAGAAGAATTTCACAAAAGTTTTAGTGGAACTAAACAAGCTGGGAAAGTTGTCGTACTCGATGAATCAATGACGTTTGATCAATTAGAAGTTGATACAGAAGTTTTAAAGCTTATCAGAGAAAACAAATCATCAACAAGAGAAATAGCAGGTGTATTTGGTATTCCATTGCATAAGTTCGGCATAGAAACAGCGAACATGAGTATCACGGATGCTAATTTAGATTACTTATCAACTTTAAAACCTTATATTACATGCGTTTGTGCAGAATTGAATTTTAAGTTTAATGATGAATATGTGAATCGTGAATTTAAATTTGATACCACTGAAATACGAGTTGTTGATGAAAAAACACAAGCTGAAATTGACAAAATTAACATTGATTCTGGAAAGATGAATATCGATGAAATTAGACAACGTGATGGATTAGCGCCAATACCAGGCGGTAATGGTAGCATTCACAGAGTCGATTTAAACCATGTAAATATTGAACTTGTAGATGAGTATCAGATGAATAAATCGAGAGCTACTGATAAAAAATTGAAAGGTGGTGAGGAAAATGAGTAAGGAAACGAGAGTTGGCAACATTATTGAGGTACGCTCAAATGATAACAACGAAATGGTCATAGAGGGGTATGCGTTAAAGTTTGACACTTGGTCTGAAAATCTTGGTGGATTCAAAGAAACGATTTCACGTCGCGCTTTAGAAAACACTGATTTATCTGATGTGCGTTGTTTAGTAGATCATATCCCATCGCAAATAATTGGTAGGACAAAATCGGGTACTTTGGAGCTCGAAACTGATGATGTTGGACTTAAATATCGTTGTAAGTTACCAAACACAACATTTGCACGTGATTTATATGAGAACATGCGTGTAGGCAACATCAATCAATGTTCGTTTGGTTTTATGCTTGACGATAAAGGCGATGAAGTGCGTTTTGATGAACAAGAAAACATTTACAAACGTACTTTAACAGCAATTCGTGAACTTACAGATGTTTCTGTAGTGACTTATCCGGCTTACAAAGACACTGATGTTAAACCAGCATTACGTAGTATTGAAACCGTTAAAAAAGAACAACGTAAAAAAGAATTAGAAATAAGACTAAAGAAACACTCTATATTAAATAATATTTGGTGAAGTTGAACACCATTATCAAATACAGCCATTGGACATGCTGAATATAGCGATGTCTATTTTTTTATGCCAATTTTAGGAGGAAATTAAATGAAAACAAAAGAAGAGTTACAATCTGAGATTTCAGACATTAAAAGACAAATTGATTTAAAGGTGAAGTATGCAACGAGAGCACTTAATAACGATGAGTTAGAAAAAGCAGAAAAATTAGAACAAGAAATTACTGATTTACGTTCTCAAATCCAAGAAAAACAAGAAGAATTAGATAAGCTAAAAGAAAAAGATGGAACTTCAGAAAACAATCAACAATCAGTGGAAGTAAACGAAGCAAGTACTTATCGAAATCAAGCAAACATTAATGATTTAGGTATTTCGATTCAAAACACAAAGGTAACATCACAAGAAGTTAGAGATTTTACTGAATATCTTGAAACACGCAATGATATTCAAGGTGGTTCGTTAAAAACAGACTCAGGATTTGTAGTTATTCCAGAGGAAATTGTTACAGATATTTTAAAATTAAAAGAGGTTGAGTTTAATCTTGATAAGTATGTGACGGTCAAACGTGTTACAAATGGTTCTGGTAAATATCCGGTAGTACGACAATCAGAAGTTGCAGCCCTTGAAAAAGTTGAAGAATTAGAAGAAAACCCTGAATTAGCAGTTAAACCATTCTTCCAATTAGCATATGACATTAATACACACCGTGGTTACTTCCGAATTTCACGTGAAGCAATCGAAGATGCAAAAGTGAATGTTTTGCAAGAATTGAAACTATGGATGGCGCGAACTATTGCAGCAACACGAAACAAAGCAATTATTGATGTTATCACTAAAGGATCAACGGGTTCTACAAGTTCAGGTTTTGAAAAAGAAGGCAAGAAATTAGAAGTTAAAAAAGCAAAATCTTTAGATGATATTAAAGATGCTATTAACCTGAATGTTAAGCCAAATTACGAACATAATGTTGCGATTGTTTCGCAAACTATGTTTGCAAAATTAGACAAAATGAAAGATAAGCTAGGAAACTATTTAATCCAGCCAGATGTTAAAGAAAAAACGCAACAGCGTTTATTAGGAGCTAAAATCGAAATTTTACCTGATGAAGTACTAGGGCAAAAAGGTAATAACACTTTGATTATCGGTAACTTAAAAGATGCGATTGTTTTATTTGACCGCTCTCAATACCAAGCATCATGGACTGACTACATGCATTTCGGAGAATGTTTAATGATTGCTGTACGTCAAGACTGTAGAATTCTAGATTATAAATCAGCAATTGTGATTGAATATGATGATAGTGAACGCGGTGAAGGCGATCTTGGCTTAGAAGCATAATAAGCGCTCGATACTTTATAAAGAGGTGATAAACTATGGCAATGTATGAAGTGAAGAAATCTTATACTGACTTGGAAAAAGGCCAGTATTTAAAGTCAGGTAAACGTGTTGAAATGACAGTAAAACGTGCTGAATATGTTAACAAAAAGCTGAAAGAGCATGGAGTAATACTTGAAAGAGTAAAAGAAGAATAGGTGATTGAATGCAATTAACAGCTGAGGAACTTAAGTTATTAAAAAAGCATTGCAAAATAGATCACAATTCAGAGGACGACTTATTAGAAATATATTACTCTTGGGCATTCCGTGAAATAGCTAGCGCTGTTACGGATAAACCAAGTAAATATATTGATTGGTTTAAAAGTCATCCTCTATTTGCTCGTGCTATATACCCTTTAGCAAGTTACTATTTTGAAAACCGTATTGCTTATTTGGATAGGGATTTATCGCTTGCGCCACATATGGTTTTAAGTACGGTGCATAAATTGAGAGGTTCATTTGAGCAATTTTTGGAGAGTGAAAATGATGAAATTTAATTCCAATAAATTAAATGAACGTATAGATTTTTGTGAAGATGTAAGCGAGAGAGTGAACGGAAATCCGATGAAACCGAAGACGAAAATATTATACTCTTGTTTCGCTTGCATTCAAGAATCTAAAGAATCCGACACTCAAACGAATCTCAATACAGGTAGCAAATTCATTAAAACTATTATTATCAGAGATACACGAGGTGATTATAAACCAACAAATAAGCATTACGTCTTGCATGAAGGGCAAAGATTTAACATCAAATATGTAAAGCCAGATTATCAAGATAAATCTTATTTGCGTATCTATGGCGAGGTGGTCATTTAATGGGGGCAAGAATTGAAAGTAATAACATCGAACAAGGTTTGAAAAATGCAGTTTTAAAAATGAATTTAAATAGTAATGTAATTGTCAAAGCTGGGGCTATGTCATTAGTCCCGCTTTTAAAAAGTAATACACCTTTTGCGAATACTAAAAAGCATGCTCGCGATCACATAGCTGTTTCTAATGTGAAAACAGACAGACACACAAGTGAGAAAATTGTTACAATTGGTTACGCTAAAGGCGTCTCACATCGTATTCATGCAACAGAATTTGGAACAATGTACCAAAAACCACAATTGTTTATAACAAAAACAGAAAAGCAAGGGAAAAACAAAGTTTTAAAAACAATGCTTGATACTGCTAAGAGGTTGCAAAAATGATTAATGTTACCAAATTAATTAGAAACGCTATTATTGCAAATAACATTACAGATGAAGTGAATGTGTTTAACTACACTATAGATGACCATTTTCACGAAAAAACTGACAAGCCTATTATTCGTATATATCCCTTACCGTTCAATCCTGACACATACGCTGATGATAACGAGATTTCAAGAGAATACCATTACCAAATTGATGTTTGGTGGTCTCAAGATGAACCGAACGAGCAAGCAGAAAAAATTGTTGAGTTACTCAAAGTGATAAATTTTCAATGTTATTACAGAGAACCGTTATACGAGAGTGACGTCATGTCATTCAGACATATTATAAGAGCAAAAGGCTCGATTTTATCAATGAAATTGGAGGAAAATTAAATGATTGAAAAATTGAAACAAGCACCAAGATTTTTAAAATTAAACTTACAACATTTTGCAGATACAGGAGTTTCGGGTATCGCAATTGGGGTATCAAACTTTTATTATGCACCTATTTTAAAAGATACAGAAAATGAATGGGAAACTGGAGCTGGCACACGTATTCGTTTCTTAAAAGAAATTGAAGTAGACCGTCCACAAGATACCGAGGAAGATTATGGGGATGATATGGTCGCAGCAACTGCTGTATCTAATGGCAAACTAAGTGTTAAGACAACATTTGTTACTGTTCCTGCTGACGATAAGGCGTTCTTGAATGGCGCTAAAAAAGGTGTAGGTGGTTATAAATATGGAGCTAAGGATATCCCGCCAGATGTAGCGATTGTATTTGAAAGACGTAATCATGATGAGTCTTCAGAATGGGTTGGCTTGTTCAAAGGTAAATTCACTCGTTCAAGCATCAAAGGGCAAACAAAACAAGATAAAGTTGAATTCCAGAATGACGACGTAGAAGGCAATTTTATTGATCGTTTGTTTGATGAGAGCTCGCATGTTACTGGCTATGATAAAAAAGGAAGCACTACAGGGCGCGATTATGTATTCATGGAAACATTTGGTAAAACTTATGATGAATTCATGTCTAGTCGAGGAGAACAAAATATGGAACCTGTAGAAAAAGAAATGAAAAAAACAGAAAAAGTTGAAGTCACTTCTGTAAACGTCACTGATGAACAAGTTACAGTTAAAGTTGATGCTACTAAACAACTATCAGCCACAACCGAACCATCTGGACAGAAAGTAACTTATGCAGTGACTGAGGGGCAAACGTATGCTAGCGTAACATCAACTGGCCTCGTTAAAGGTTTGGCGGAAGGTAATGCGACCGTTACAGCGACTGCAGGAAAGCAAACTGATACTGTGCAAATTACAGTACAATCTAATTTAGAAATGTAAGTTTTGAGGGCTTAACGCCCTCTTTTTATTTTGGCCAAATTAAAAAGAAAGTAGGAATTTAATAATGGAACGTACATCAATTGAATTAATTACAGGATTTACAAAAACAGGAAAGCCGCAATATCAAAAGTATTTAGCGAAGCCGATTATTACTTTGTTTGAAACAATTCAAGGTTCAAAATTAGGTTTGAAACTTAACAAAGCCTTTAAGGGGGCTGATTTTAAAGATCTAACAGAAGAAGAATTTAATAACTTAAGTGTGACAGAACAGGAAGAATACAAAAACAAGCAAGAAGAATACGAAAACAACATGGCTGTACAAATGGAAGTATTAGAAGAAGTTTTGGATTTCATCGTTGAAGCTTTTGATAATCAATTTACCAGTATAGAACTTCAAAAAGGATTACCAAATGGTCAAGAAGGTATTGAAAAGATTGGACAGTTAATTGGACGAATTACAGGTGGGGAACCTAGCGATACAAAAAAGTTCGTGACAGAGAATCAGAAATAAGAAAAGAAGATTTAACACCTGAAGCTGTCTACAACAATTACAGGAAAATAGCTAAAGATTTGATAGAAAAAGGCATGGATGCAGAAAAAGTGGCTAACATGCCGATACACTTCTTTTTAGACATTGTCGAATCGAAGATTGAAACAAAGCGAACTGCGAAAAGTTTTAAAGATATTTTTTAATCAGCCTTTAAAGGTTGATTTTTTATTTACATCTTGGAAGAAAGGAGGTTTTTAAATGCCTAATCCTATAGGTAATATGGTCATAAAGGTTGATTTAGATGGTTCTGGATTCAATAGAGGTGTGACAGGTTTAAATAGGCAAATGAAAATGGTTTCGCGTGAGCTTTCGGCTAATTTATCACAATTTTCTAGATATGATAATTCATTAGAAAAGTCGAAGATAAAAGTCGAAGGTTTGAGTAAAAAACAAAAAGTTCAAGCCCAGATTACTAAAGAGCTGAAAGATAGTTATGACAAACTTAGTAAAGAAACTGGTGAAAACAGTGCAAAGACACAAGCTGCGGCTGCTAAATACAATGAAGCTTACGCTAAATTAAACCAATATGAGCGAGAGTTAAATCAAGCCACACAAGAATTAAAAGACATGCAAAGAGAGCAGAAAGCATTAAATACTGCAATGGGAAAACTTGGTACCAACTTTAATAATTTTGGTCCTAAACTTCAAGAAATTGGTAACAGTATGAAAAATGTAGGCCGTAACATGACTATGTATGTAACTGCGCCGGTGGTTGCTGGGTTTGCTGTAGCAGCTAAAAAAGGTATTGAATTCGATGACAGTATGAGAAAAGTTAAAGCAACTTCAGGTGCTACTGGGGAAGAGTTTGAAGCTTTGAAGAAAAAGGCTCGCGAAATGGGTGCAACAACAAAATTTAGTGCATCAGATTCGGCTGAAGCATTAAATTACATGGCACTTGCTGGTTGGGATTCTAAGCAAATGATGGAAGGTTTAAGCGGAGTTATGGATTTAGCGGCAGCATCTGGCGAAGAACTGGAAGCAGTAAGTGACATTGTTACAGATGGACTAACGGCATTCGGTTTAAAAGCAAAGGATAGTGGTCATTTTGCGGACATTTTAGCACAAACTAGCTCGAAGGCAAATACGGATGTTAGAGGGCTCGGAGAAGCTTTTAAATATGTCGCTCCTGTAGCAGGTGCGTTAGGTTACACGATTGAAGATACATCTATTGCGATAGGTTTAATGAGTAATGCTGGTATCAAAGGTGAAAAAGCAGGTACAGCGTTACGAACAATGTTCACCAATCTTTCAAGTCCAACTAGAGCTATGGGGAATGAAATGGAACGCTTAGGAATATCTATTACAGATAGTAATGGGAAAATGATTCCTATGCGAAAGCTTTTAGACCAACTGAGGGAAAAATTTAAACATCTTTCAAAAGACCAACAAGCTAGTTCTGCAGCTACAATATTTGGTAAAGAAGCGATGTCAGGAGCATTAGCGATTATAAATGCTTCTGATGAAGACTATCAAAAGTTAACCAAATCTATAGATTCATCTACCGGCGCATCTAAAAGAATGGCCGATACAATGGAATCTGGTTTAGGTGGGAAATTAAGAACTTTAAGGTCGCAATTAGAAGAACTAGCCTTAACGATTTATGACAGAATAGAACCAGCACTAAAGATTATAGTAAGTGCTTTTAGCAAAGTAGTGACATGGGTTACTAAATTACCAACGTCAATTCAATTAGCGGTTGTTGGGTTTGGATTATTTGCAGCAGTTTTAGGTCCTTTAGTTTTTATGTTCGGTTTATTTATCAGCGTGATGGGGAATGCAATGACAGTTTTAGGACCCTTGTTAATAAACGTTAATAAAGCTGGTGGTTTATTCGCGTTTTTAAGAACTAAAATCGCATCACTTGTTAAACTATTTCCGATTTTAGGTGTGTCGATATCAAGTTTAACGTTACCTATAACATTAATTGTAGGTGCATTAGTTGGTATTGGCATAGCTTTCTATCAAGCTTATAAACGTTCAGAAACTTTTAGAAATATTGTAAATCAGGCAATCTCTGGTGTAGCAAACGCATTTAAAGCAGCTAAACTAGCGTTACAAGGTTTCTTTGATTTATTCAAAGGTGATAGTAAAGGCGCGGTTACCCTAGAGAAGATATTTCCACCCGAAACTGTAGCAGGAATACAAAATGTAGTTAATACGATTAGAACAACTTTCTTTAAAGTAGTTGATGCAATCGTTGGTTTCGCCAAAGAGATAGGCGCTCAATTAGCCTCTTTCTGGAAAGAGAACGGCTCAGAAATAACACAAGCTTTGCAAAATATAGCTGGTTTCATTAAAGCAACCTTTGAATTTATTTTTAACTTTATTATTAAACCAATCATGTTTGCGATTTGGCAAGTGATGCAATTTATTTGGCCGGCGGTTAAAGCTTTGATTGTCAGCACTTGGGAAAATATCAAAGGTGTAATACAAGGGGCTATTAATATTATTTTGGGTATTATCAAAGTGTTCTCTAGTCTTTTCACAGGAAACTGGCGAGGCGTTTGGGACGGCATTGTAATGATACTGAAAGGTACTGTGCAGTTAATTTGGAATTTAATACAACTGTGGTTTGTAGGTAAGATTCTAGGTGTTGTTAGATACTTTGGTGGATTGCTTAAAGGTTTAATATCCGGTATCTGGGGTGTTATCAAAGGTATTTTCACAAAATCATTATCTGCAATTTGGAATGCAACGAAAAGTATTTTTGGTTTCTTATACAATAGTGTTAAATCTATTTTCACTAATATGAAAAACTGGTTATCTAGTACGTGGAATAATATCAAAAGCAATACCGTCGGCAAGGCTCATTCGTTATTTACGGGTGTAAGGTCTAAATTCACAAGTTTATGGAATGCGACGAAAGATATATTTACTAAATTAAGAAATTGGATGTCAAACATCTGGAACTCTATTAAAGATAACACGGTAGGTATAGCTGGTCGTTTGTGGGATAAAGTACGTAATATCTTCGGAAACATGCGTGACGGTTTAAAATCTATCATTGGTAAAATTAAAGATCATATCGGCGGTATGGTAGATGCTATTAAAAAAGGACTTAATAAATTAATTGAAGGCTTAAACTGGGTCGGTGGTAAGTTAGGTATGGATGAAATACCTAGGTTACACACTGGTACAGAGCACACACATACTACTACAAGATTAGTTAAGAACGGTAAGATTGCACGTGATACATTCGCTACAGTTGGGGATAAAGGACGTGGAAATGGTCCAAATGGTTTTAGAAATGAAATGATTGAATTCCCTAATGGTAAACGTGTAATCACACCTAGTACAGACACTACTGCTTATTTACCTAAAGGCTCAAAAGTATACAACGGTGCACAAACTTATTCAATGTTAAACGGAACGCTTCCGAGATTTCATTTCGGTACTACTATGTGGAAAGATATTAAATCTAGTGCATCATCGGCATTTAACTGGACAAAAGATCAAATAGGTAAAGGCACAAAGTGGCTTGGCGATAAAGTTGGTGATGTCATGGACTTTATCGATAATCCAGGCAAACTTTTAAATTATGTACTTCAAGCGTTTGGAGTTGATTTCAGTTCTCTAACTAAAGGTATGGGTATTGCTGGCGATATAACAAAAGCTGCATGGTCTAAGATTAAGAAAAGTGCAATCAAGTGGCTTGAGGATGCTTTCGCAGAGTCGGGTGATGGCGGTGTATTAGATATGAGTAAATTACGTTACTTATACGGTCACACTGCTGCTTATACACGAGAAACCGGACGCCCATTCCATGAAGGTCTGGATTTTGATTACATTTACGAACCTGTTCCATCAACCATTAATGGTAGAGCACAAGTTATGCCTGTTCATAATGGTGGTTATGGAAAATGGGTGAAAATTGTAAAGGGCGCCTTAGAAGTTATTTATGCACATTTATCTAAATATAAAGTTAAAACTGGTCAACAAGTTAGGGTCGGACAGACTGTTGGTATATCGGGGAATACGGGGTTTAGTACAGGACCTCACTTACATTATGAGATGCGTTGGAATGGAAGACATAGAGACCCGTTACCGTGGTTAAGAAAGAATAATGGGGGCGGCAAAAGTACACCCGGTGGTAATGGTGCAGCTAATGCTAGACGAGCTATTAAGGCTGCTCAAAATATTTTAGGAGGAAGGTATAAGGCGAGTTGGATTACTAACGAGATGATGCGTGTTGCGAGTCGTGAATCCAATTATACAGCTAATGCAGTCAATAATTGGGATAGCAACGCAAGAGCTGGTATACCTTCAAGAGGTATGTTCCAAATGATAGATCCTTCATTTAGAGCGTACGCAAAGTCGGGTTACAATAATCCTCTCAACCCAACTCATCAAGCTATATCGGCTATGAGATATATTGTGGGTAAATGGGTACCAAGAACAGGCTCATGGAGAGCTGCGTTCAAACGCGCTGGTGATTACGCATATGCTACTGGTGGCAAAGTCTATAACGGATTGTATCACTTAGGGGAAGAAGGATATCCAGAGTGGATAATACCTACTGATCCAAGTAGAGCGAACGAAGCACACAAATTATTAGCTTTAGCTGCTAACGATATTGATAACCGCTCTAAAAATAAGCGACCAAACAACTTACCAAATCCAAGTATAAGTAATAGTGATACAAACTATATTCATACATTGGAGAATAAACTGGATGCGGTTATTAATTGTTTGGTTAGTTTGGTTGAGTCTAATCAAGTTATTGCAGATAAGGATTACGAACCAGTTATTAATAAGTATGTGTTTGAAGATGAGGTAAATAATTCTATCGATAAACGAGAGCGTCACGAATCTACAAGAGTTAGATTTAGAAGAGGAGGCACGATAATCTAATGCAAGATACAATTCAAATAGACAATAAAACAATTGGATGGCTGGTTGTGCAAAGAGGGTTCGAGATACCCTCTTTTAATTTTGTTACTGAAAAAGAAAACGTAAAAGGTAGAGCGGGATCTATTGTTAAGAATCGTTATTTAAATGATATCGAATTTGATTTACCATTAATTATTCGAAACGAAAAATTGTCACCAGGTGGAGAAAAAACACACGATGATATATTAGAAGCATTGGTCAAGTTCTTCAATATTAAAGATTTAACACCTAAAAAACTTAAATTCAAATCTCAAAACTGGTATTGGTTTGCATATTTTGATGGTCCATTAAAATTACCGAAAAACCCAAGAGGTTCAGTGAAGTTCACTATAAAAGTAGTGTTAACAGATCCTTATAAATACTCGGTAACTGGAAACAAAAACACCGCGATTTCAGACCAAGTTTCAGTTGTAAATAGTGGGACTGCTGACACTCCTTTAATTGTTGAAGCCCGAGCAATTAAACCATCTAGTTACTTTATGATCACTAAAAATGATGAAGATTATTTTATGGTTGGTGATGATGAGGTAACCAAAGAAGTTAAGGATTACATGCCTCCTGTTTATCATAGTGAGTTTCGTGATTTCAAAGGTTGGACTAAGATGATTACTGAAGATATTCCAAGTAATGATTTAGGTGGTAAGGTCGGCGGTGACTTTGTGATATCCAATCTTGGCGAAGGATATAAAGCAACTAATTTTCCTGATGCAAAAGGTTGGGTTGGTGCTGGCACGAAACGAGGGCTCCCTAAAGCGATGACAGATTTTCAAATTACCTATAAATGTATTGTTGAACAAAAAGGTAAAGGTGCCGGAAGAACAGCACAACATATTTATGATAGTGATGGTAAGTTACTTGCTTCTATTGGTTATGAAAATAAATATCATGATAGAAAAATAGGACATATTGTTGTTACGTTGTATAACCAAAAAGGAGACCCCAAAAAGATATACGACTATCAGAATAAACCGATAATGTATAACTTGGACAGAATCGTTGTTTATATGCGGCTCAGAAGAGTAGGTAATAAATTTTCTATTAAAACTTGGAAATTTGATCACATTAAAGACCCAGATAGACGTAAACCTATTGATATGGATGAGAAAGAGTGGATAGATGGCGGTAAGTTTTATCAGCGTCCAGCTTCTATCATAGCTATCTATAGTGCGAAGTATAACGGTTATAAGTGGATGGAGATGAATGGATTAGGTTCATTCAATACGGAGATTCTACCGAAACCGAAAGGCGCAAGGGATGTCATTATACAAAAAGGTGATTTAGTGAAAATAGATATGCAAGCAAAAAGTGTTGTCATCAATGAGGAACCAATGTTGAGCGAGAAATCGTTTGGAAGTAATTATTTCAATGTTGATTCTGGGTACAGTGAATTAATCATACAACCTGAAAACGTCTTTGATACGACGGTTAAATGGCAAGATAGATATTTATAGAAAGGAGATGAGAGTGTGATACATGTTTTAGATTTTAACGACAAGATTATAGATTTCCTTTCTACTGATGACCCTTCCTTAGTTAGAGCGATTCATAAACGTAATGTTAATGACAATTCAGAAATGCTTGAACTGCTCATATCATCAGAAAGAGCTGAAAAGTTCCGTGAACGACATCGTGTTATTATAAGGGATTCAAACAAACAATGGCGTGAATTTATTATTAACTGGGTTCAAGATACGATGGACGGCTACACAGAGATAGAATGTATAGCGTCTTATCTTGCTGATATAACAACAGCTAAACCGTATGCACCAGGCAAATTTGAGAAAAAGACAACTTCAGAAGCATTGAAAGATGTGTTGAGCGATACAGGTTGGGAAGTTTCTGAACAAACCGAATACGATGGCTTACGTACTACGTCATGGACTTCTTATCAAACTAGATATGAAGTTTTAAAGCAATTATGTACAACCTATAAAATGGCATTGGATTTTTATATAGAGCTTAGTTCTAATACCGTCAAAGGTAGATATGTGGTACTCAAAAAGAAAAACAGCTTATTCAAAGGTAAAGAAATTGAGTATGGTAAAGATTTGGTTGGGTTAACTAGGAAGATTGATATGTCAGAAATCAAAACAGCATTAATTGCTGTGGGACCCGAAAATGACAAAGGAAAGCGTTTAGAGTTAGTTGTGACTGATGACGAAGCACAAAGTCAATTCAACTTACCTACCCGTTATATTTGGGGAATATACGAACCTCAATCAGATGATCAAAATATGAATGAAACACGGTTGCGTTCTTTAGCCAAAACAGAGTTAAATAAACGTAAGTCGGCAGTTATGTCATATGAGATTACTTCTACTGATTTGGAAGTTACGTATCCGCACGAGATTATATCAATTGGTGATACAGTCAGAGTAAAACATAGAGATTTTAACCCGCCATTGTATGTAGAGGCAGAAGTTATTGCCGAAGAATATAACATAATTTCAGAAAATAGCACATATACATTCGGTCAACCTAAAGAGTTCAAAGAATCAGAATTACGAGAAGAGTTTAACAAACGATTGAACATAATACATCAAAAGTTAAACGATAATATTAGCAATATCAACACTATAGTTAAAGATGTTGTAGATGGTGAATTAGAATACTTTGAACGCAAAATACACAAAAATGATACACCGCCAGAAAATCCAGTCAATGATATGCTTTGGTATGATACAAGTAACCCTGATGTTGCTGTCTTGCGTAGATATTGGAATGGTCGATGGATTGAAGCAACACCAAATGATGTTGAAAAATTAGGTGGTATAACAAGAGAGAAAGCGCTATTCAGTGAATTAAACAATATTTTTATTAATTTATCTATACAACACGCTAGTCTTTTGTCAGAAGCTACAGAATTACTGAATAGCGAGTACTTAGTAGATAATGATTTGAAAGCGGACTTACAAGCAAGTTTAGACGCTGTGATTGATGTTTATAATCAAATTAAAAATAATTTAGAATCTATGACACCCGAAACTGCAACGATTGGTCGGTTGGTAGATACACAAGCTTTATTTCTTGAATATAGAAAGAAATTACAAGATGTCTATACAGATGTAGAAGATGTCAAAATCGCTATTTCAGATAGATTTAAATTATTACAGTCACAATACACTGATGAAAAATATAAAGAAGCGTTGGAAATAATAGCAACAAAATTTGGTTTAACGGTGAATGAAGATTTGCAGTTAGTCGGAGAACCTAATGTTGTTAAATCAGCTATTGAAGCAGCTAGAGAATCCACAAAAGAACAATTACGTGACTATGTAAAAACATCGGACTATAAAACAGACAAAGACGGTATTGTTGAACGTTTAGATACTGCTGAAGCTGAGAGAACGACTTTAAAAGGTGAAATCAAAGATAAAGTTACGTTAAACGAATATCGAAACGGATTGGAAGAACAAAAACAATATACTGATGACCAGTTAAGTGATTTGTCCAATAATCCTGAGATTAAAGCAAGTATTGAACAAGCAAATCAAGAAGCGCAAGAAGCTTTAAAATCATACATTGATGCTCAAGATAATCTTAAAGAGAAGGAATCGCAAGCGTATGCTGATGGTAAAATTTCGGAAGAAGAGCAACGCGCTATACAAGATGCTCAAGCTAAACTTGAAGAGGCAAAACAAAACGCAGAACTAAAGGCTAGAAACGCTGAAAAGAAAGCTAATGCTTATACAGACAACAAGGTCAAAGAAAGCACAGATGCACAGAGGAGAACACTGACTCGCTATGGTTCTCAAATTATACAAAATGGTAAGGAAATCAAATTAAGAACTACTAAAGAAGAGTTTAATGCAACCAATCGTACACTTTCAAATATATTAAACGAGATTGTCCAAAACGTTACAGATGGAACAACAATCAGATATGATGATAACGGAGTGGCTCAAGCTTTAAATGTGGGGCCACGTGGTATTAGATTAAATGCTGATAAAATTGATATTAACGGTAATAGAGAAATAAACCTTCTTATCCAAAATATGCGAGATAAAGTAGATAAAACCGATATTGTCAACAGCCTTAATTTATCAAGAGAGGGTCTTGATATCAATGTTAATAGAATTGGAATTAAAGGCGGTAACAATAACAGATATGTTCAAATACAGAATGATTCTATTGAACTAGGTGGTATTGTGCAACGAACTTGGAAAGGCAAACGATCAACCGATGATATATTCACACGTCTTAAAGATGGACATCTAAGGTTTAGAAATAATACCGCAGGCGGTTCACTTTATATGTCACATTTTGGTATTTCAACATATATTGATGGAGAAGGCGAAGACGGAGGTTCATCCGGTACTATTCAATGGTGGGATAAAACTTACAGTGATAGCGGTATGAATGGCATAACAATCAATTCCTATGGTGGTGTCGTTGCACTAACGTCAGATAATAATCGGGTTGTTCTGGAGTCTTACGCTTCATCGAATATCAAAAGCAAACAGGCACCGGTGTATTTATATCCAAACACAGACAAAGTGCCTGGATTAAACCGATTTGCATTCACGCTGTCTAATGCAGATAACGCTTATTCGAGTGATGGTTATATTATGTTTGGTTCTGATGAGAACTATGATTACGGTGCGGGTATCAGGTTTTCTAAAGAAAGAAATAAAGGTCTTGTTCAAATTGTTAATGGACGATATGCAACAGGTGGAGATACAACAATCGAAGCAGGGTATGGCAAATTTAATATGCTGAAACGACGTGATGGTAATAGGTATATTCATATACAGAGTACAGACCTACTGTCTGTAGGTTCAGATGATGCAGGAGATAGGATAGCTTCTAACTCAATTTATAGACGTACTTATTCGGCCGCAGCTAATTTGCATATTACTTCTGCTGGCACAATTGGGCGTTCGACATCAGCGCGTAAATACAAGTTATCTATCGAAAATCAATATAACGATAGAGATGAACAACTGGAACATTCAAAAGCTATTCTTAACTTACCTATTAGAACGTGGTTTGATAAAGCTGAGTCTGAAATTTTAGCTAGAGAGCTGAGAGAAGATAGAAAATTATCGGAAGACACCTATAAACTTGATAGATACGTAGGTTTGATTGCTGAAGAGGTGGAGAATTTAGGATTAAAAGAGTTTGTCACGTATGATGACAAAGGAGAAATTGAAGGTATAGCGTATGATCGTCTATGGATTCATCTTATCCCTGTTATCAAAGAACAACAACTAAGAATCAAGAAATTGGAGGAGTCAAAGAATGCAGGATAACAAACAAGGATTACAAGCTAATCCTGAATATACAATTCATTATTTATCACAGGAAATTATGAGGTTAACACAAGAAAACGCGATGTTAAAAGCGTATATACAAGAAAATAAAGAAAATCAACAATGTGCTGAGGAAGAGTAATCCTTAGCACTATTTTTATACAAAAATTTAAGGAGGTCATTTAATTATGGCAAAAGAAATTATCAACAATACAGAAAGGTTTATTTTAGTACAAATCGACAAAGAAGGTACAGAACGTGTAGTATATCAAGATTTCACAGGAAGTTTTACAACTTCTGAAATGGTTAACCATGCTCAAGATTTTAAATCTGAAGAAAACGCTAAGAAAATTGCGGAGACGTTAAATTTGTTATATCAATTAACTAACAAAAAACAACGTGTGAAAGTAGTTAAAGAAGTAGTTGAAAGATCAGATTTATCTCCAGAGGTAACAGTTAACACTGAAACAGTATGAAAAGCTATGAGTTAGATACTCATAATCTTTATTCTTTTAGAAAGCGGGTGTACTGAATTGGGGTGGTTCAAAAAACACGAACATGAATGGCGCATCAGAAGGTTAGAAGAGAATGATAAAACAATGCTCAGCACACTCAACGAAATTAAATTAGGTCAAAAAACCCAAGAGCAAGTTAACATTAAATTAGATAAAACCTTAGATGCTATTCAAAAAGAAAGAGAAATAGATGAAAAGAATAAGAAAGAAAATGATAAGAACATACGTGATATGAAAATGTGGGTGCTTGGTTTAGTTGGGACAATATTTGGGTCGCTAATTATAGCATTATTGCGTATGCTTATGGGCATATAAGAGAGGTGAATAAAATGTTTAAACTAATCTTTGGTTATAGTTTCTGGACATGTTTTTGGTTCGGTAAATGTAAATAAGTTTTAGTCAGTGCTTCGGTACTGACTTTTTATTTATTGTTGTAATTATGGTAATATGCAGAAGTGAGCAAGTTGGATAGATGGTGGCTATCTGAGTATAAGGAGGTGGTGCCTATGGTGGCATTACTGAAATCTTTAGAAAGGAGACGCCTAATGATTACAATTAGTACCATGTTGCAGTTTGGTTTATTCCTTATTGCATTGATAGGTCTAGTAATCAAGCTTATTGAATTAAGCAATAAAAAATAACCATCGCTAACTTTGGCTGGTTTCGATGGTTAAATGGTTATTAATTTAATCTTTAATCTAAAATAGCCACCGTCTTTTTAACGGGCTCATTAGGGTAACATGTTTGCGCATGTTGCCCTTTTTCTATATATAAATTAACACACCATAATATAAATATCAAATAGACGGCTTATTAGTCGTCTTTTTATTTTGGGTAAAAGGAGATAAGAATATGATTAATTGGAAAATTAGAATGAAACAAAAATCATTTTGGGTAGCGATATTGTCAGCTATCTTTTTATTTGCTCAAAACATCGCAAAAGCTATTGGGTATGATATCCAAGTTTATACAGAGCAATTAACAGACGGTTTAAACGCTATATTAGGATTTTTAGTATTAACTGGTGTGATTCAAGACCCGACTACTAAAGGTATAGGTGATAGCCACCAAGCTTTAGAATATGAAGAACCAAGAAGAAAATACTAGGAGGTAAAATAATGAAAACATACAGTGAAGCAAGAGCAAGGTTACGTTGGTATCAAGGTAGATATATTGATTTTGACGGTTGGTATGGTTACCAATGTGCAGATTTAGCAGTTGATTACATTTATTGGTTGTTAGAAATTAGAATGTGGGGAAATGCAAAAGATGCAATCAATAACGATTTTAAAAACATGGCAACAGTATATGAAAACACACCATCGTTTGTTCCACAAATAGGTGATGTGGCTGTATTTACCAAAGGAATATATAAACAATACGGTCATATTGGTTTAGTGTTTAATGGTGGTAATACAAACCAATTTTTAATTTTGGAACAGAACTATGACGGTAACGCAAATACGCCTGCAAAGTTACGTTGGGATAATTATTACGGCTGTACTCACTTTATTAGACCTAAGTATAAAAGTGAGGGCTTAATGAATAAGATCACAAATAAAGTTAAACCACCTGCTCAAAAAGCAGTCGGTAAATCTGCAAGTAAAATAACAGTTGGAAGTAAAGCGCCTTATAACCTTAAATGGTCAAAAGGTGCTTATTTTAATGCGAAAATCGACGGCTTAGGTGCTACTTCAGCCACTAGATACGGTGATAATCGTACTAACTATAGATTCGATGTTGGACAGGCTGTATACGCGCCTGGAACATTAATATATGTGTTTGAAATTATAGATGGTTGGTGTCGCATTTATTGGAACAATCATAATGAGTGGATATGGCATGAGAGATTGATTGTGAAAGAAGTGTTTTAATTCTTAGGTTAAAATGTTAAATATTTGTTAATTATTTTTTAATGTAAGTTTAGTTTCTTTTAATATTTTATTGATTTTTAATATTTTTTCGATATAAAATGAAGTTGTTGATATTTATCATCTTAAATAAGGGTGTTAGCTATAAAAAGAGATAAATAAAAACAAATATATTATATTTGGAGGAAGCGCCATGCTCAAAAGAAGTTTATTATTTTTAACTGTTTTATTGTTATTATTCTCATTTTCTTCAATTACTAATGAGGTAAGTGCATCAAGTTCATTCGACAAAGGAAAATATAAAAAAGGCGATGACGCGAGTTATTTTGAACCAACAGGCCCGTATTTGATGGTAAATGTGACTGGAGTTGATGGTAAAGGAAATGAATTGCTATCCCCTCATTATGTCGAGTTTCCTATTAAACCTGGGACTACACTTACAAAAGAAAAAATTGAATACTATGTCGAATGGGCATTAGATGCGACAGCATATAAAGAGTTTAGAGTAGTTGAATTAGATCCAAGCGCAAAGATCGAAGTCACTTATTATGATAAGAATAAGAAAAAAGAAGAAACGAAGTCTTTCCCTATAACAGAAAAAGGTTTTGTTGTCCCAGATTTATCAGAGCATATTAAAAACCCTGGATTCAACTTAATTACAAAGGTTATTATAGAAAAGAAATAAAACAAAATAGTTGTTTATTATAGAAAGCAATGTCTTGATTGAATATGTGTAGTGAAAATTATCTTTCATCAAATTCTCATTCATGCACGAATGGTTCTTCCCCACCTAATCAGATATTAGGTGACTTATGGGGAGAAATCAGTTAGGATGAAAAAGTGGATAATCCTTTTTTAGGCAGGTACTTCGGTACTTGCCTATTTTTTTATGTTATAATCTTTCTAGACGTATTCAAGGGACGTCTTTTTAGATTGTATGTTATAGCTAGCTTTCGGGCTAGTTTTTTGTTATGATGTGTTACACATGCATCAACTATTTACATCTATCCTTGTTCACCCAAGCATGTCACTGGGTGTTTTTTCTTATGATAGAGAGCATAGTTTTCATACTACTCCCTCGTAGTATATATGACTTTAGCATTCCCGTATAATAGTTTACGGGGTGCTTTTTATGTTATAATTAACTGTATATAGTAGGAGTGAACTATATAGCCTGTTAAGTGGCCTAGTAACCTAACACTTATCCTGCAATTGATATCCTTTTTGCCCTTCACTCGATACATATATCTCAACAACATAGAAATATTACAGTCGCTACACCGCATCTTAAATGGTGTGGTTATTTTTATTGGAAGTGTGTATCAGGTATCAGTAATGTTAAAACACCAGCTAAAAATGAAAAGAATTCACCAGTGCCAGCAGGTTATACACTCGATAAAAACAATGTACCGTATAAAAAAGAGACTGGTTATTACACAGTTGCCAATGTTAAAGGTAATAACGTGAGGGATGGCTATTCAACTAATTCAAGAATTACAGGTGTATTACCCAATAACGCAACTATCAAATATGACGGCGCATATTGCATTAATGGCTATAGATGGATTACTTATATTGCTAATAGTGGACAACGTCGTTATATAGCGACAGGAGAGGTAGACAAGGCAGGTAATAGAATAAGCAGTTTTGGTAAGTTTAGTGCAGTTTGATAATTAGATATATAAAGGTTTGGCAAGTTATGAAATGTCTGCCAAACCTTTATATAAAAAAGAAATATCTACCTTTTAATCCGAGGTATGAAAACGAGAATTGGACCTTTACAGAATTACTCTATGAAGCGCCATATTTAAAAAGCTACCAAGACGAAGAGGATGAAGAGGATGAGGAGGCAGATTGCCTTGAATATATTGACAATACTGATAAGATAATATATCTTTTATATAGAAGATATCGCCGTATGTAAGGATTTCAGGGGGCAAGGCATAGGCAGCGCGCTTATCAATATATCTATAGAATGGGCAAAGCATAAAAACTTGCATGGACTAATGCTTGAAACCCAGGACAATAACCTTATAGCTTGTAAATTCTATCATAATTGTGGTTTCAAAATCGGCTCCGTCGATACTATGTTATACGCCAACTTTCAAAACAACTTTGAAAAAGCTGTTTTCTGGTATTTAAGGTTTTAGAATGCAAGGAACAGTGAATTGGAGTTCGTCTTGTTATAATTAGCTTCTTGGGGTATCTTTAAATACTGTAGAAAAGAGGAAGGAAATAATAAATGGCTAAAATGAGAATATCACCGGAATTGAAAAAACTGATCGAAAAATACCGCTGCGTAAAAGATACGGAAGGAATGTCTCCTGCTAAGGTATATAAGCTGGTGGGAGAAAATGAAAACCTATATTTAAAAATGACGGACAGCCGGTATAAAGGGACCACCTATGATGTGGAACGGGAAAAGGACATGATGCTATGGCTGGAAGGAAAGCTGCCTGTTCCAAAGGTCCTGCACTTTGAACGGCATGATGGCTGGAGCAATCTGCTCATGAGTGAGGCCGATGGCGTCCTTTGCTCGGAAGAGTATGAAGATGAACAAAGCCCTGAAAAGATTATCGAGCTGTATGCGGAGTGCATCAGGCTCTTTCACTCCATCGACATATCGGATTGTCCCTATACGAATAGCTTAGACAGCCGCTTAGCCGAATTGGATTACTTACTGAATAACGATCTGGCCGATGTGGATTGCGAAAACTGGGAAGAAGACACTCCATTTAAAGATCCGCGCGAGCTGTATGATTTTTTAAAGACGGAAAAGCCCGAAGAGGAACTTGTCTTTTCCCACGGCGACCTGGGAGACAGCAACATCTTTGTGAAAGATGGCAAAGTAAGTGGCTTTATTGATCTTGGGAGAAGCGGCAGGGCGGACAAGTGGTATGACATTGCCTTCTGCGTCCGGTCGATCAGGGAGGATATCGGGGAAGAACAGTATGTCGAGCTATTTTTTGACTTACTGGGGATCAAGCCTGATTGGGAGAAAATAAAATATTATATTTTACTGGATGAATTGTTTTAGTACCTAGATTTAGATGTCTAAAAAGCTTTAACTACAAGCTTTTTAGACATCTAATCTTTTCTGAAGTACATCCGCAACTGTCCATACTCTGATGTTTTATATCTTTTCTAAAAGTTCGCTAGATAGGGGTCCCAAAAGATTTATAACGAAATTGACGAAGCACTAAAAAGTAAATATTAAAAAAACCACCCTTTTACGGGTGGTTTTAATTTTCTAGATAATATAAAAGTGTTCATAAATAAAACAGTATAGGCAAACAATAAAGTATTGAAAAAAGTAAGTTTAATATGAAAATTGTTAAATGAACGACATCTTTTGTTTTTATAAATATCAAGAAAATAATCAAACTCAAAATAAATAACGTAACTGTAGTCATAGGCGTCCATACATAATCAGCATTAGTCATTAAGAATGGTGCAGCCATTATGAAAAAATTTATAATGCAGATGAAATAGACAATTAGACTATAAATTAGGTAAATAACAATACACACCCTTCATAAATAAATAATTTAAATCCTATATATTTTAACAAAAGTAAAACACAGAAGTGTAGAAAATAAAAAATATTGGTAAATAAAATCAATAAGTTTAACCAATATGTTGCTCGCTTCATACCGTATATTGCAACAAAAATTCCGATCAAGAAAAATATAGCCCCTATGATAAAACAGAAATCCGATGCTGAACTATTAAAAAATGAGGTGTTTAGAGTTAGAAAATGAGTTAATGAGTTGACTATAACTAATAAGATATTAATTATATTTGTATGGTTCTTCACATGATACCTCCAAGTAAAAAAATCTAATTAATAAAGTGAATGCTTGATGAACAAGCAGTTATTCCAAACAGAATCAATAAGAAAAGTAGAATCAACATGCTAATGCCCCATAAACAACCCTTTTCACTTTCTCTATTATTAATTTCTTGACTTCTTTTTAAAGATTTATTACTTTTACATTCTTTAGTTGTTTTAAATTTCACGTTTTTATTACTTCCTTTTGTCTAAAAGTTTACAATGAATTTTTGATTATAATAATATATTCAAAATAGTACTATCTAGTTTGATATGTCAAGCAATATTATTATAAAATTGGAATTCTGAGTTGTCTACTCTAATTTATTATATTTACCTATAAAAATACACCTCAAAAAATAGATTTTTCAGTCTAGCTTTTGGGGTGTACATTCCACACAAACATGTGATTATTTTGATGTTTCTATTAAACTTGTAATTTTAAATTTAAAGTCCCTAAAAAGTCCCTAAAATTTTATTTTATATGGGGTATTATTGATAATGATAAAGTTATAAACCTTGATATTATGCTGTTTTACTTTTTGAATGATAAGTAATTTTATGTTAAAAGTCT**

**Supplementary tables**

**Table S1: Strains**

| Strain | Description | Reference/ Origin |
| --- | --- | --- |
| *Escherichia coli* |  |  |
| DC10B |  | (1) |
| *Staphylococcus aureus* |  |  |
| 8325-4 (RN0450) | NCTC8325 cured of Φ11, Φ12 and Φ13 | (2) |
| 8325-4 Φ13K | Single-lysogen, *kan^R^* | (3) |
| 8325-4 Φ13K-*rep* | Single-lysogen, carrying replication deficient phage mutant (3), *kan^R^* | (3) |
| SH1000 | *rsbU* repaired derivative of 8325-4 | (4)  Susanne Engelmann, TU Braunschweig, Germany |
| SH1000 Φ13K | Single-lysogen, *kan^R^* | (3) |
| SH1000 Φ13K-*rep* | Single-lysogen, carrying replication deficient phage mutant (3), *kan^R^* | (5) |
| Newman-c | Phage-cured | (6) |
| Newman-c Φ13K | Single-lysogen, *kan^R^* | (3) |
| Newman-c Φ13K-*rep* | Single-lysogen, carrying replication deficient phage mutant (3), *kan^R^* | This study |
| MW2c | Phage-cured | (7) |
| MW2c Φ13K | Single-lysogen, *kan^R^* | (3) |
| MW2c Φ13K-*rep* | Single-lysogen, carrying replication deficient phage mutant (3), *kan^R^* | (3) |
| SH1000 Φ13K-*TATA* | Single-lysogen, *p23* TATA-Box substitution, *kan^R^* | (5) |
| Newman-c Φ13K-*TATA* | Single-lysogen, *p23* TATA-Box substitution, *kan^R^* | (5) |
| SH1000 Φ13K-*ltr* | Single-lysogen, carrying *ltr* deficient phage mutant, *kan^R^* | This study |
| Newman-c Φ13K-*ltr* | Single-lysogen, carrying *ltr* deficient phage mutant, *kan^R^* | This study |
| RN4220-331 | *ΔmazEFrsbUVWsigB*::*tetM* | (8) |
| 8325-4 Φ13K *sigB* | Single-lysogen, *sigB*::*tetM* | This study |
| SH1000 Φ13K *sigB* | Single-lysogen, *sigB*::*tetM* | This study |
| Newman-c Φ13K *sigB* | Single-lysogen, *sigB*::*tetM* | This study |
| SM2 | Newman *spoVG*::*erm*, *yabJ-spoVG* mutant, *erm^R^* | (9)  Markus Bischoff |
| SH1000 Φ13K *spoVG* | Single-lysogen, *spoVG*::*erm* | This study |
| LS1 |  | (10)  Löffler, Münster, Germany |
| RN4220 | restriction deficient derivate of 8325-4, rK-mK+ | (11) |

**Table S2: Oligonucleotides**

| Oligonucleotide | Sequence | Used for |
| --- | --- | --- |
| pCG896gibfor | gctggcggccgctgcatgGGATCAT  GAGCATTCTTGATATAGGC | Cloning pCG896 |
| pCG896gibrev | cataaataatcatcctcctaagCCCTC  ACTTAATGTGAGAGTTCA | Cloning pCG896 |
| pcIyfpoutsidecontrolfor | GGACAGGTATCCGGTAAGCG | Cloning |
| pcIyfpcontrolrev | TGACAAGTGTTGGCCATGGA | Cloning |
| pCG896insidecontrolfor | GGACACATCGTACAGTTCGG | Cloning |
| pCG910SDMfor | cggccGCTGTGTAGCAAAACATTTA  TATTTC | Cloning pCG910, pCG925 |
| pCG910SDMrev | cggcgAAAAACAATATGTAGCATCA  AAATTAG | Cloning pCG910, pCG925 |
| pCG943SDMfor | TAATTTTGATGCTACATATTGTTTT  TTATTATAATTG | Cloning pCG943 |
| pCG943SDMrev | CGGTTTCTTGTTGCAAG | Cloning pCG943 |
| pIMAYcontrolfor | CCAGCCCCCTCACTACAT | Cloning pCG925, pCG926 |
| pIMAYcontrolrev | ATCACCCGACGCACTTTG | Cloning pCG925, pCG926 |
| pCG925gibfor | aattcctgcagcccggggCTATGACTATT  GTATTTGCTATATTGCT | Cloning pCG925 |
| pCG925gibrev | gccgctctagaactagtgGCACATCACTC  CTTGTCGAC | Cloning pCG925 |
| pCG925outsidecontrolfor | GCGGAGGTAAGTGAGTGA | Cloning pCG925 |
| pCG925outsidecontrolrev | GGATGACCACATCGCTTCA | Cloning pCG925 |
| pCG926insert1gibfor | aattcctgcagcccggggAGACATC  TTAGATCGAGTTAAGGAGG | Cloning pCG926 |
| pCG926insert1gibrev | ttcctgtttTACATGCAATACCT  CCGATA | Cloning pCG926 |
| pCG926insert2gibfor | ttgcatgtaAAACAGGAAAGAAA  TACGTGA | Cloning pCG926 |
| pCG926insert2gibrev | gccgctctagaactagtgAATAGACA  ATGCACATCACTCCT | Cloning pCG926 |
| 926outsidecontrolfor | CGACCAACTCATTGACGC | Cloning pCG926 |
| 926outsidecontrolrev | AACCATAGTCGCTTGATTGCCACA | Cloning pCG926 |
| circlefor | TTTTATTTTATATGGGGTATTATTGA | qPCR (Φ13) |
| circlerev | GTGTATTCTCATTTGTTAGAAGAAAA | qPCR (Φ13) |
| SAOUHSC_02200qPCRfor | GGCACGACTAGCAATAAA | RT-qPCR (*ltr*) |
| SAOUHSC_02200qPCRrev | GTCTCTGCCTATATCAAGAAT | RT-qPCR (*ltr*) |
| cIqPCRfor | AGAACGTCAAGATGAAACGA | RT-qPCR (*cI*) |
| cIqPCRrev | AATTCTTCTCCTATGCCAGC | RT-qPCR (*cI*) |
| SAOUHSC02234DIGfor | TAATACGACTCACTATAGGGAG  ATGCAAAATTGTACTGAGTGC | RT-qPCR (*mor*) |
| SAOUHSC02234DIGrev | ATGTGTTACGACTACTCACG | RT-qPCR (*mor*) |
| SAOUHSC02196for2 | CACGAATCAAAACGGCATTA | RT-qPCR (*terL*) |
| SAOUHSC02196rev2 | ACAACAATCGAATCAATGGC | RT-qPCR (*terL*) |
| SAOUHSC02191for | TTTGCATCTTCGATTGCTTC | RT-qPCR (*mcp*) |
| SAOUHSC02191DIGrev | TACGACAATCAGAAGTTGCA | RT-qPCR (*mcp*) |
| RTqPCRamidasefor | AAATAGGTGATGTGGCTGTA | RT-qPCR (*amidase*) |
| RTqPCRamidaserev | AGTGAGTACAGCCGTAATAATT | RT-qPCR (*amidase*) |
| holin255for | ATGATTAATTGGAAAATTAGAA | RT-qPCR (*holin*) |
| holin255rev | CTAGTATTTTCTTCTTGGTTCT | RT-qPCR (*holin*) |
| sakLClo | CATCAAGTTCATTCGACAAAGGAAA | RT-qPCR (*sak*) |
| sak-A | TGTAGTCCCAGGTTTAATAGG | RT-qPCR (*sak*) |
| asp493f | AAAATTGCTGGTATCGCTGC | RT-qPCR (*asp*) |
| asp848r | TGTAAACCTTGTCTTTCTTGGT | RT-qPCR (*asp*) |
| T7-SAOUHSC02196DIGfor | TAATACGACTCACTATAGGGAG  ATTAGGGTCTGGAAGCATTTC | DIG-probe Northern Blot (*terL*) |
| SAOUHSC02196DIGrev | ATGGTTGCCATTGGGATAAT | DIG-probe Northern Blot (*terL*) |
| T7-SAOUHSC02191DIGfor | TAATACGACTCACTATAGGGA  GATTTGCATCTTCGATTGCTTC | DIG-probe Northern Blot (*mcp*) |
| SAOUHSC02191DIGrev | TACGACAATCAGAAGTTGCA | DIG-probe Northern Blot (*mcp*) |

**Table S3: Plasmids**

| Plasmid | Description | Resistance casette | Reference/ Origin |
| --- | --- | --- | --- |
| pIMAY-Z | Mutagenesis vector | *cm* | (12) |
| pCG725 | P*_cap_*-*yfp* | *cm* | (8) |
| pCG896 | Promoter construct P_23_-*yfp* | *cm* | This study |
| pCG910 | Promoter construct P_23_-*TATA*-*yfp* | *cm* | This study |
| pCG925 | Mutagenesis vector for p23 TATA Box mutation (pIMAY-Z) | *cm* | (5) |
| pCG926 | Mutagenesis vector for *ltr* mutation (pIMAY-Z) | *cm* | This study |
| pCG943 | Promoter construct P_23_-Δ*repeat*-*yfp* | *cm* | This study |
